# The equilibrium between two quaternary assembly states determines the activity of SPOP and its cancer mutants

**DOI:** 10.1101/2025.06.19.659812

**Authors:** Matthew J. Cuneo, Ömer Güllülü, Mohamed-Raafet Ammar, Xinrui Gui, Kelly Churion, Martin Turk, Brian G. O’Flynn, Nafiseh Sabri, Tanja Mittag

## Abstract

Proteostasis is critical for preventing oncogenesis. Both activating and inactivating mutations in the ubiquitin ligase subunit SPOP result in oncogenesis in different tissues. SPOP assembles into filaments that are multivalent for substrates, and substrates have multiple weak motifs for SPOP that are not activated via post-translational modifications. It is thus unclear how regulation is achieved. Here, we show that SPOP filaments circularize into rings that dimerize into up to 2.5 MDa-large, auto-inhibited double donuts. The equilibrium between double donuts and linear filaments determines SPOP activity. Activating and deactivating cancer mutations shift the equilibrium towards the filament or the double donut, respectively, and this influences substrate turnover and subcellular localization. This regulatory mechanism requires long filaments that can circularize into rings, likely explaining the presence of multiple weak SPOP-binding motifs in substrates. Activating and deactivating mutations combine to give rise to intermediate activities, suggesting new levers for cancer therapies.

**Highlights:**

- SPOP assemblies exist in an equilibrium between circular double donuts and linear filaments.
- Double donuts occlude access to the substrate binding site and are inactive.
- Mutations in different cancers shift the equilibrium towards active or inactive states.
- Regulation through these structural transitions requires large filamentous assemblies.

## Introduction

Ubiquitin ligases play critical roles in the maintenance of cellular proteostasis by flagging proteins for proteasomal degradation, thereby keeping protein levels in check and maintaining cellular health. Given the destructive effects of ubiquitin ligase activity, it requires tight regulation. Indeed, recurrent mutations in ubiquitin ligases such as BRCA1, VHL, FBW7, MDM2 and SPOP are common drivers of oncogenesis because they lead to dysregulation of the levels of relevant substrates ^1,2^. Some ubiquitin ligases achieve regulation by recognizing only post-translationally modified substrates. For example, VHL, a substrate receptor of the Cullin-2-RING-ligase (CRL2), recognizes hydroxylated HIF-α and thereby reads out oxygen levels in kidney tissues ^3-6^. Cdc4, a substrate receptor of the Cullin-1-RING-ligase (CRL1), only recognizes substrates that are phosphorylated by cyclin-dependent kinases in specific phases of the cell cycle, marking them as cell cycle phase—specific substrates ^7-9^. On the other hand, other ubiquitin ligases, such as the Speckle–type POZ protein (SPOP), recognize unmodified linear motifs in their substrates ^10,11^, which raises the question of how substrate recognition is regulated. That regulation is important is underlined by the observation that both activating and deactivating mutations in *SPOP* result in cancer, albeit in different tissues. Many well-understood oncogenic SPOP mutations, including those in prostate cancer, are loss-of-function mutations that reduce or abrogate substrate binding ^12,13^. By contrast, mutations found in endometrial cancer patients activate SPOP ^14^. SPOP is also regarded an important tumor suppressor across cancers ^2,15^.

We previously showed that SPOP, which functions as substrate receptor in the Cullin-3-RING ligase (CRL3), assembles into helical filaments via its two dimerization domains, BTB and BACK ^16,17^, and additional contributions from the substrate-binding MATH domain ^17^ (**Fig. 1A,B**). Each SPOP monomer within the filament can interact via its MATH domain with a SPOP-binding motif in a substrate ^10,18,19^ (**Fig. 1C**), often with affinities in the hundreds of micromolar to millimolar range ^19,20^. Multiple SPOP-binding motifs in a single substrate can interact with several monomers in a SPOP filament and lead to tight binding and specific recognition ^19,20^.

**Figure 1.**
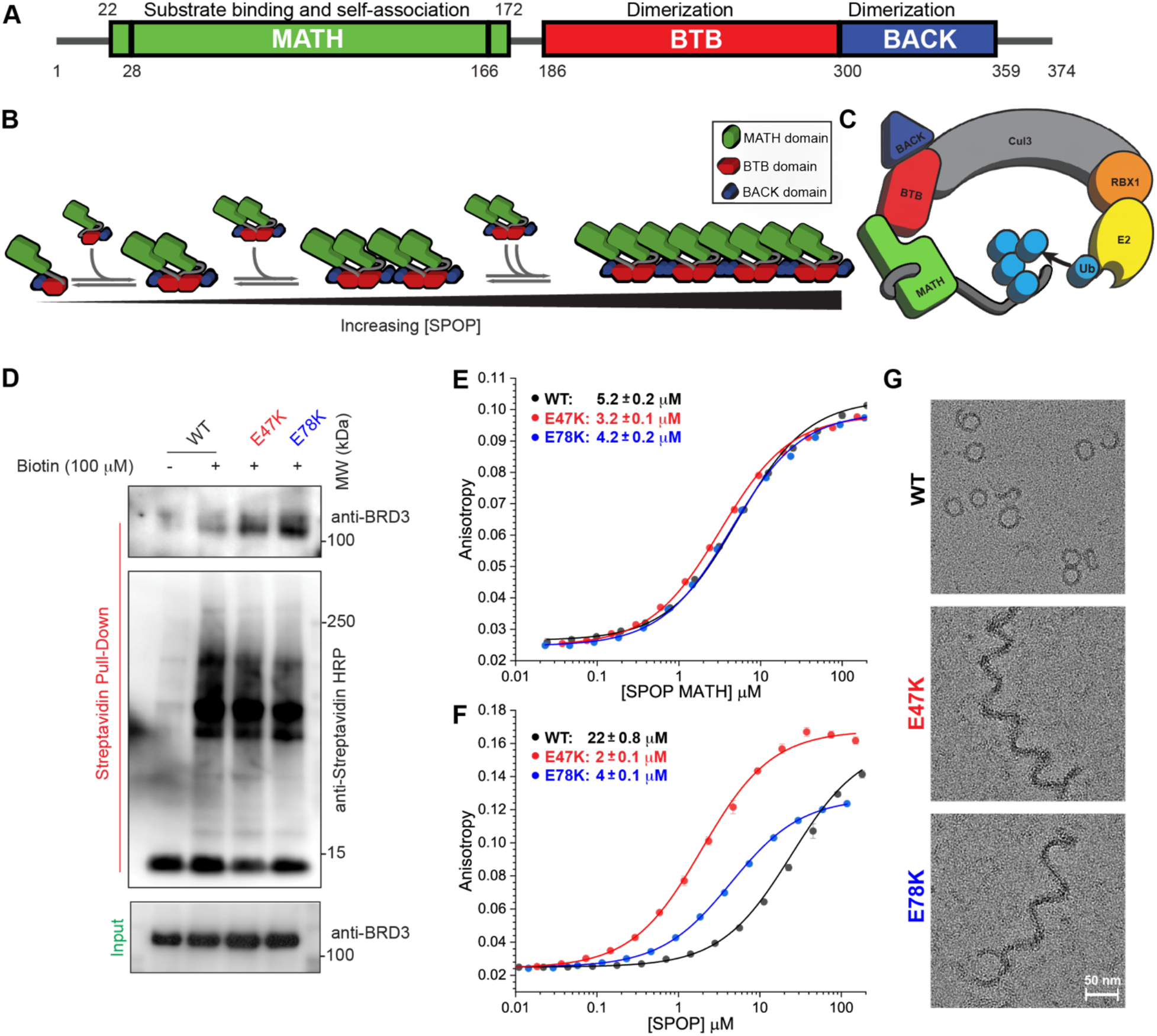
Endometrial cancer-associated mutations activate SPOP substrate binding. **(A)** Box plot of the domain architecture of SPOP. Each of the three domains are involved in oligomerization. **(B)** Cartoon schematic of the self-association behavior of SPOP. The BTB and BACK domains of SPOP each dimerize. The MATH domains engage head to tail with adjacent MATH domains. **(C)** Schematic of the CRL3^SPOP^ complex as a monomer. Cul3: Cullin-3; RBX1: RING-box protein 1; E2: ubiquitin-conjugating enzyme; Ub: ubiquitin. **(D)** Expression of TurboID-SPOP variants and subsequent pull-down of biotinylated proteins demonstrate that wild-type (WT) SPOP minimally interacts with BRD3 in mammalian cells, whereas more BRD3 interacts with SPOP E47K and E78K (14 ± 2-fold and 18 ± 4-fold increase (mean of three replicates ± the S.E.M.) compared to WT, respectively). **(E, F)** Fluorescence anisotropy binding curves of a BRD3 peptide binding to (E) the MATH domain of SPOP or to (F) full-length SPOP assemblies. Activating endometrial cancer-associated mutations do not affect MATH domain binding but enhance full-length SPOP binding up to ten-fold. **(G)** Negative–stain TEM images show WT SPOP forms donut-shaped assemblies, whereas endometrial cancer-associated mutants form linear filaments.

Previous work showed that endometrial cancer–associated mutants of SPOP are hyperactive towards a set of substrates including BRD2, BRD3 and BRD4 ^14^, although the mutant MATH domains do not harbor increased affinities for the substrates ^21^. This is rationalized by the fact that the mutated residues are located outside the substrate-binding groove of the MATH domain instead of targeting substrate-interacting residues. Our recent cryo-EM structures of the oligomeric state of wild-type (WT) SPOP and mutants revealed that the mutant W22R formed an assembly with altered quaternary structure ^17^, providing an explanation for its hyperactivity. In contrast, the structure of the endometrial mutant E47K was identical within experimental error to that of the WT, failing to explain the gain-of-function phenotype of mutation E47K and many others. Moreover, SPOP mutations are sporadically found in other cancers, but if they do not affect the substrate-binding groove, it is not known whether they mediate loss or gain of function.

It is unclear why SPOP–substrate interactions are split into many weak interactions, instead of a single strong interaction, as is more typical for other ubiquitin ligase substrate receptors. Oligomerization is a hallmark of allosteric regulatability of metabolic enzymes, and their transitions into filament states can change the conformations of active sites and regulate activities in response to conditions such as starvation-induced pH changes ^22^ or the presence of allosteric regulators ^23^. Could oligomerization regulate the activity of ubiquitin ligases as well? Does the filamentous architecture of SPOP enable regulation of its activity, and does this explain the effects of mutations found in cancer patients?

Here we address these questions and examine the molecular underpinnings of SPOP activation. We assess different gain- and loss-of-function SPOP mutations found in patients with endometrial and other types of cancer. We optimized conditions for WT SPOP cryo-EM samples to mitigate dissociation at the air–water interface and discovered that WT SPOP assembles into complexes of up to 2.5 MDa and with a double donut architecture. Through a combination of structural, biophysical, biochemical and cell biological analyses, we show that SPOP double donuts are inherently inactive and exist in an equilibrium with the active filament state. Binding of Cullin-3 (Cul3) shifts the equilibrium to the filament state and activates SPOP. Activating cancer mutations bypass the inactive double–donut state, whereas several inactivating cancer mutations stabilize the double donut instead of inactivating the substrate binding site, pointing towards possible new therapeutic strategies. Notably, the differences between the filament and double–donut states are not subtle conformational changes that directly alter the activity but rather large-scale quaternary transitions that result in steric hindrance of substrate binding. This work has implications for understanding the regulation of large ubiquitin ligase complexes and other enzymes.

## Results

### Endometrial cancer-associated mutations enhance SPOP activity via altered quaternary assembly

Endometrial cancer-associated mutations in SPOP, such as E47K and E78K, lead to hyperactivation of SPOP characterized by enhanced turnover of certain substrates such as BRD3^14^. However, how these mutations enhance substrate turnover remains unclear. To test the effects of this mutation on the interaction between SPOP and BRD3 in cells, we performed proximity ligation of SPOP interactors, expressing SPOP WT, E47K or E78K fused to the TurboID enzyme. We detected strong interactions of SPOP E47K and E78K with BRD3, whereas SPOP WT showed much weaker interaction (**Fig. 1D**).

We then examined the in vitro interactions of the isolated MATH domain and full-length SPOP with a peptide containing the main SPOP-binding motif of BRD3. In the isolated MATH domain, the E47K and E78K mutations did not result in enhanced binding to the BRD3 peptide compared to the WT (**Fig. 1E**). In contrast, full-length SPOP E47K and E78K bound the BRD3 peptide with up to ten-fold higher affinity than full-length SPOP WT (**Fig. 1F**). These data provide the first evidence that gain of function of endometrial cancer mutants is linked to enhanced binding to a substrate. Thus, the individual WT, E47K and E78K MATH domains do not possess different affinities for BRD3, but full-length SPOP WT and mutants do.

We previously showed that SPOP can form higher order assemblies and thus asked if those assemblies could account for the different binding activities of WT and mutant. We visualized the assemblies formed by WT E47K and E78K SPOP by negative-stain EM (nsEM): While E47K and E78K formed the long filaments we had previously described ^17^, WT SPOP mostly formed rings that coexisted with some filaments (**Fig. 1G**).

To understand whether different quaternary assembly states were specific for certain mutants or explanatory of a more general pattern, we visualized additional cancer-associated mutants. Endometrial cancer-associated mutants like R45W, E47A, E50K and R99W appeared as filamentous assemblies. R121Q and D140N showed mixtures of filaments and rings (**Fig. 2A**). Prostate cancer-associated mutants such as Y87F, D130E, W131G, and W131L also formed rings. Only M117V and K129E formed assemblies of different types that we did not characterize further (**Fig. 2A**). In addition, we characterized two mutants from other cancer types. E47Q, identified in a patient with adrenocortical carcinoma, formed filaments. Even an activating mutation outside of the MATH domain, R221C, found in a patient with bladder urothelial carcinoma, adopted mixtures of filaments and rings (**Fig. 2A**).

**Figure 2.**
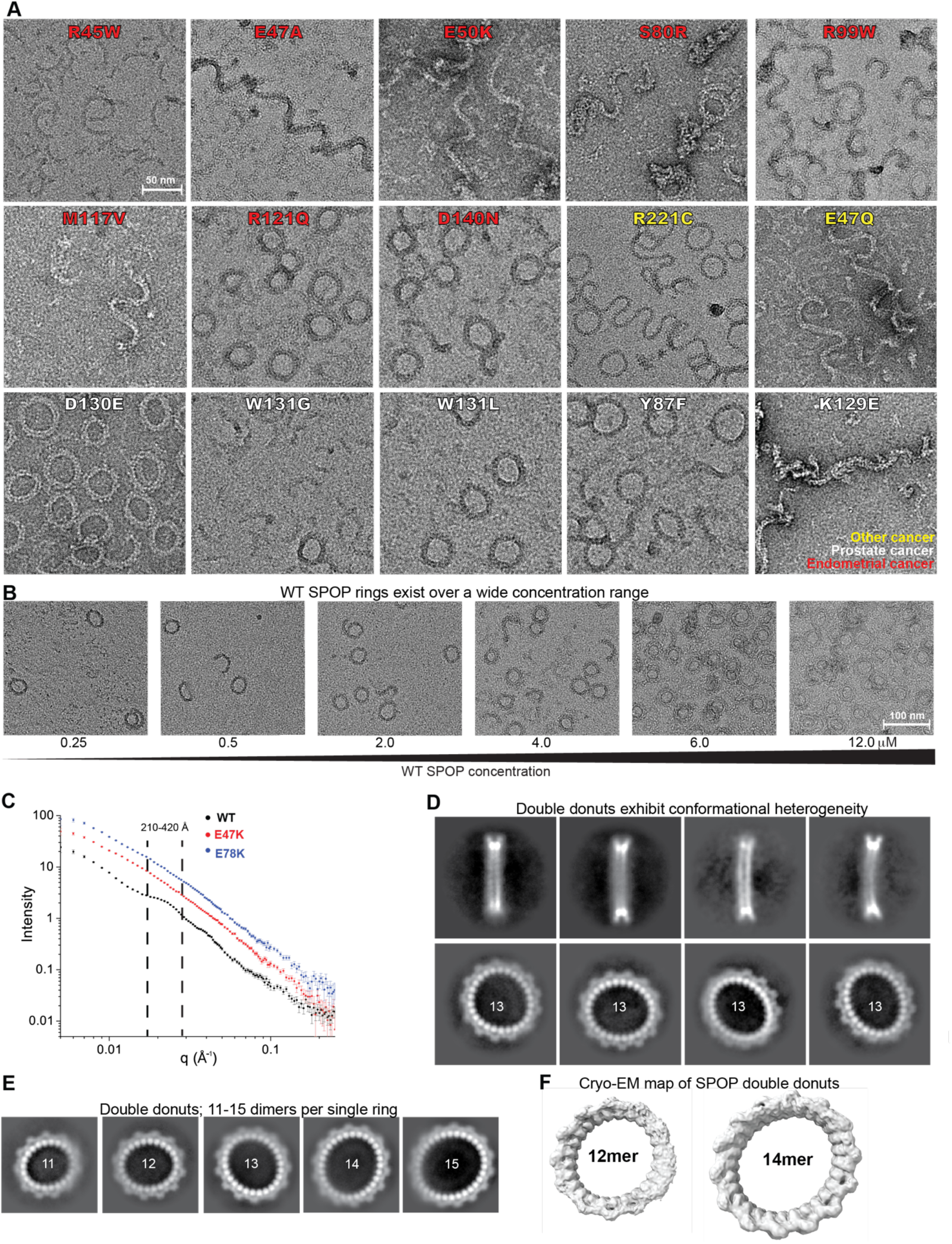
Activating mutations disrupt the ring state of WT SPOP and of inactivated mutants. **(A)** Negative–stain TEM images of a selection of SPOP mutants show activating mutants (found in patients with endometrial cancer and other cancers) adopt mostly the linear filamentous state, deactivating prostate cancer-associated mutants adopt mostly a ring state. K129E is an exception and adopts a linear multiple–filament form. **(B)** Negative–stain TEM images show that WT SPOP rings persist over a wide concentration range. **(C)** SAXS data show correlation peaks for WT SPOP consistent with distances found in rings. In samples of SPOP E47K and E78K, correlation peaks are not observable. **(D)** 2D classification of particles in WT SPOP cryo-EM samples. Top, side-views of SPOP rings show a double layer suggesting single rings dimerize to form double donuts. Top and bottom, the donuts exhibit significant conformational diversity. **(E)** 2D classes show double donut sizes range from 11–15 dimers per single ring. **(F)** Cryo-EM maps of a 12-mer (i.e., 12 dimers per single ring, giving rise to a 2.0 MDa complex formed by 48 monomers) and a 14-mer (i.e., 14 dimers per single ring giving rise to a 2.3 MDa complex formed by 56 monomers) SPOP double donut.

### SPOP ring assemblies block access to the substrate binding sites

Our observations suggested a potentially general pattern, in which WT SPOP and inactivating mutants, such as those driving prostate cancer pathogenesis, favored a ring assembly state, whereas activating mutants adopted the previously described filament state ^17^. We hence characterized the newly identified SPOP rings further.

The rings had a narrow size distribution and were stable over a wide concentration range (**Fig. 2B**). Small-angle X-ray scattering (SAXS) analysis showed the presence of correlation peaks in data from WT SPOP samples (**Fig. 2C**). The most pronounced correlation peak, centered at a q-value of 0.02 Å^-1^, is consistent with distances across the ring, with the broadness of the peak encompassing scattering vectors ranging from ∼210-420 Å, and a second smaller distance, which we could not assign to any features of the rings without additional structural information. These data indicate the presence of SPOP rings in solution. The correlation peaks were absent in data from E47K and E78K SPOP, indicating the lack of scattering vectors arising from such ring features in solutions of endometrial mutants (**Fig. 2C**).

To obtain higher-resolution information, we performed cryo-EM analysis of SPOP WT assemblies. We noticed that vitrification interfered with the ring assemblies, which are unstable at air-water interfaces. Mild cross-linking with glutaraldehyde allowed us to maintain and visualize rings (**Fig. S1A, S2A**). Initial 2D classification of the assemblies revealed the characteristic patterns created by the BTB and BACK oligomerization domains on the ring exterior. Side views revealed that the ring assemblies have two layers, resembling two stacked donuts, and will be referred to as “double donuts” henceforth. Distances between the two rings likely give rise to the correlation peak indicative of a smaller distance in the SAXS data (**Fig. 2C**). Based on the characteristic patterns on the exterior, the double donuts associate via their MATH domain surfaces. Side views also revealed that the double donuts are not planar but accommodate a variety of different out-of-plane deformations (**Fig. 2D**). Further classification revealed distinct sizes of double donuts ranging from 11 SPOP dimers per single ring (with a diameter of ∼360 Å) to up to 15 dimers (and a diameter of ∼520 Å) (**Fig. 2E**). We did not observe any single rings. Classification of double donut sections showed the presence of distinct in-phase and out-of-phase forms, in which the BTB and BACK domains of the two rings were aligned in phase and out of phase relative to each other (**Fig. S1B**). Their separate refinement resulted in 3D densities at resolutions of 4.3 Å (**Fig. S1C,D**) and showed that the arrangement of domains in a single ring is highly similar to those in the filaments we have previously visualized ^17^.

Treating the double donuts as single particles, we refined three-dimensional classes that contained 12, 13, or 14 dimers per single ring (**Fig. 2F and Fig. S2A-C**). Our maps converged at resolutions of ∼10 Å (**Fig. S2D**), and the out-of-plane dynamics of the double donuts (**Fig. 2D**) hindered our ability to superimpose the rings *en bloc* and to improve the resolution further. SPOP thus assembles into complexes ranging from 1.8–2.4 MDa, with holes in the donut rings large enough to accommodate individual ribosome particles (**Fig. S2E**).

To investigate the function of these large assemblies, we sought to visualize the donut-donut interface at higher resolution. Optimization of freezing conditions allowed us to maintain and visualize a mixture of linear filaments and rings in the absence of cross-linker (**Figure S3A**). In both in-phase and out-of-phase configurations of the double donuts, all MATH domains can be aligned, enabling focused refinement of six MATH domains at a time, three from each donut, to visualize the donut-donut interface (**Fig. S3B**). Our final maps had resolutions of 3.2 Å (**Fig. S3C**) and enabled modeling of the protein side chains into the densities (**Fig. 3A,B** and **Fig. S3D**). The interface bears many residues that are mutated in endometrial, prostate and other cancer types (**Fig. 3A**). The substrate binding grooves of the MATH domains are buried in the interface (**Fig. 3B**), indicating that the double donut may be an inactive state of SPOP.

**Figure 3.**
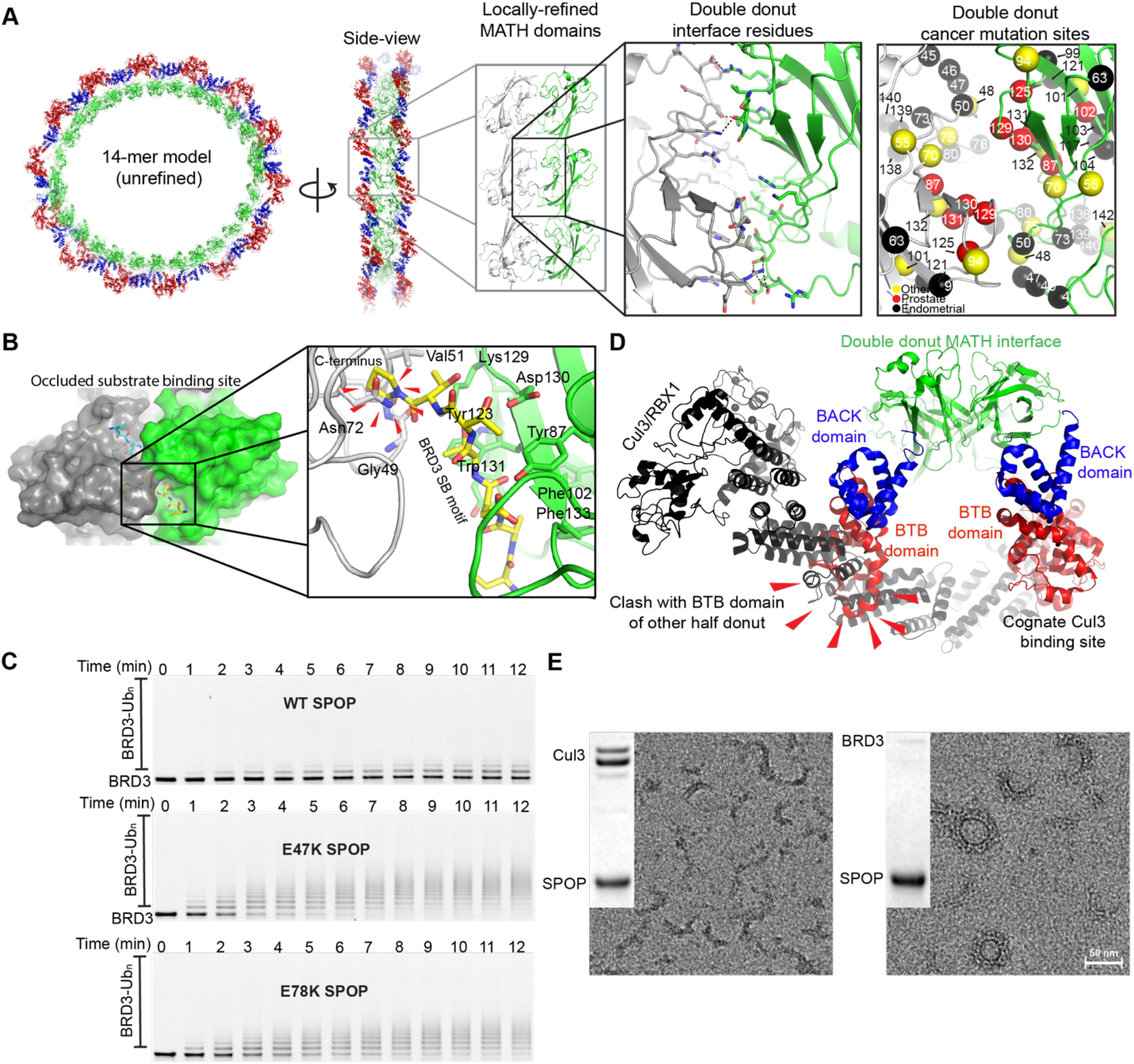
Cryo-EM structure of the SPOP double donut reveals occluded substrate- and obstructed Cul3-binding sites. **(A)** Model of a 14-mer in-phase double donut created through superposition of partial donut assemblies. Local refinement of the MATH domain portion of partial donut assemblies (gray box) enabled reconstruction of the donut–donut interface (black box). The donut–donut interface is lined with mutations found in cancer patients (red box). **(B)** The substrate-binding groove is buried in the donut–donut interface. Superposition of a MATH domain bound to a BRD3 peptide (yellow and cyan carbon atoms for the peptides on both sides of the interface, pdb 6I41) shows significant clashes with the opposite MATH domain (red arrows). **(C)** Time course of in vitro ubiquitination assay of WT SPOP or activating mutants ubiquitinating BRD3 protein. **(D)** Superposition of the Cul3/RBX1 complex onto one ring in the double donut shows large clashes with the BTB domain of the second ring. **(E)** SPOP was mixed at 1:1 ratio with Cul3 (left, partially neddylated) or BRD3 (right), and complexes were purified by size-exclusion chromatography and visualized by negative–stain TEM. BRD3 binds SPOP with sub-stoichiometric ratio.

To test this hypothesis, we performed in vitro ubiquitination assays with purified components to reconstitute neddylated CRL3^SPOP^ and using BRD3 as the substrate. WT SPOP was considerably less active than mutants E47K and E78K (**Fig. 3C**), showing that the cellular activity differences are recapitulated by the reconstituted complexes and must therefore be mediated by intrinsic differences in properties. Nevertheless, WT SPOP showed a moderate level of activity, leading us to ask whether the double–donut state, with its buried substrate binding grooves, supports any enzymatic activity. We modeled SPOP double donuts in complex with Cul3 and found substantial steric clashes (**Fig. 3D**). However, we were able to generate SPOP–Cul3 complexes in vitro and separate them from small species by size-exclusion chromatography (SEC). nsEM analysis revealed assemblies that seemed to be Cul3-decorated SPOP filaments, with a notable absence of donuts (**Fig. 3E**). We reasoned that substrates should also stabilize the filament state, yet the SEC–isolated fractions containing SPOP and BRD3 analyzed by nsEM contained a mixture of curved filaments as well as ring shapes (**Fig. 3E**). These data suggest that substrate binds to a minor population of SPOP, i.e. the filaments. Thus, we wondered whether the equilibrium between double donut and linear filament dictates SPOP activity.

### The double–donut state can be further stabilized than in WT SPOP

We hypothesized that SPOP WT assemblies exist in an equilibrium between presumably inactive double donuts and active filaments, that substrate can only bind to the filamentous portion of the assemblies, but that Cul3 binding initiates dissociation of double donuts into filaments. To test this hypothesis, we sought to generate SPOP mutants that stabilize the double donut form. We mutated key residues in the MATH domain involved in the donut-donut interface (**Fig. 4A**). We then analyzed the assemblies formed by the SPOP mutants by nsEM. We found that the V51E, D77E, S96R, and G128D mutants preserved the double–donut character of SPOP (**Fig. 4B**), whereas others (G49E, L76R, Q127E, K129Y) adopted different assemblies and possibly aggregated states (**Fig. S4**). We analyzed the four mutants that formed double donuts for their ability to bind and ubiquitinate BRD3. With the exception of S96R, the mutants showed reduced in vitro ubiquitination activity towards BRD3 compared to WT (**Fig. 4C**) and reduced binding affinities to an SB-motif containing peptide (**Fig. 4D)**. However, when these mutations were introduced into the isolated MATH domain, the mutants bound the BRD3 peptide with similar affinity as the WT MATH domain (**Fig. 4D**), showing that the mutations did not compromise the substrate-binding groove. Thus, reduction in substrate binding in the mutants was not caused by loss of binding affinity of the MATH domain, but rather by the inaccessibility of the substrate binding site in the double–donut state. Overall, we identified three SPOP mutants that stabilized the double–donut state and reduced activity to different extents. Given the similar activities of G128D and V51E, we restricted our following investigations to D77E and V51E.

**Figure 4.**
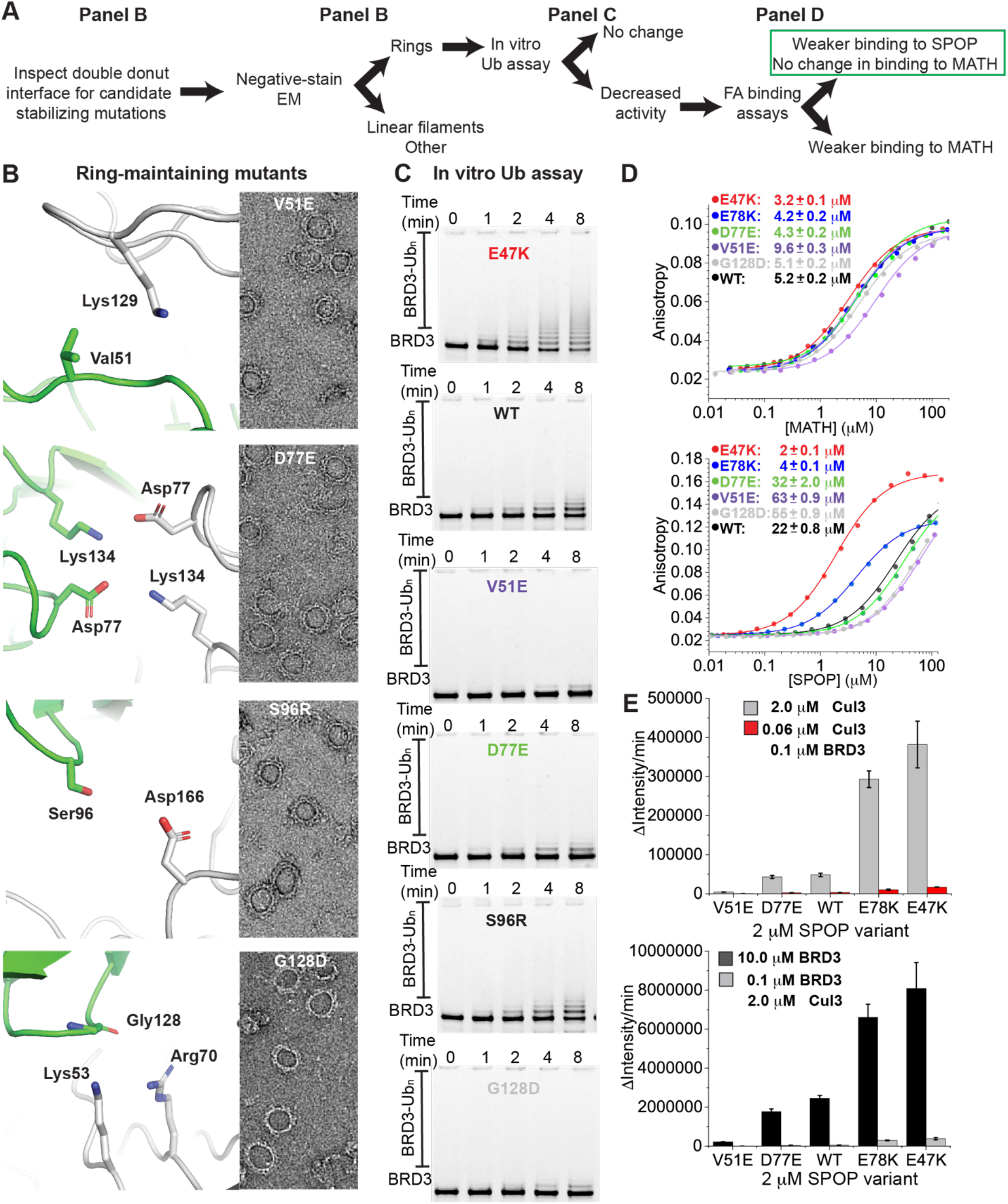
Rational design of double donut–stabilizing mutants of SPOP shows that the equilibrium with the filamentous form determines SPOP activity. **(A)** A decision tree for the experimental process of designing double donut– stabilizing mutants of SPOP. **(B)** The high-resolution structure of the MATH assemblies across the donut–donut interface was used to identify candidate mutations (left panels). The mutants were subjected to negative–stain TEM (right panels) to identify those mutants that did not disrupt the ring state. **(C)** In vitro ubiquitination assays were carried out to identify mutants that were less active than WT SPOP. E47K and WT are shown for comparison. **(D)** Fluorescence anisotropy binding experiments with BRD3 peptide to quantify binding to either the MATH domain (top) or full-length SPOP (bottom) WT and mutants. The values in the legend represent *K*_D_ values as the mean from triplicate experiments with the error as the S.E.M. Wild-type data is the same as in Fig. 1E. **(E)** Initial velocities from in vitro ubiquitination assays with BRD3 protein. Top, assays were carried out at fixed BRD3 protein concentration (0.1 μM) at either low (0.06 μM) or stoichiometric Cul3 (2 μM) concentration relative to SPOP (2 μM). Bottom, assays were carried out at fixed Cul3 concentration (2 μM) at either low (0.1 μM) or high BRD3 (10 μM) concentration. Gray bars in top and bottom plots are the same data. Initial velocities are mean values from at least triplicate experiments with the error bar representing the S.D.

### The equilibrium between inactive double–donut and active filament states determines SPOP activity

Next, we used the designed mutants to test the hypothesis that endometrial cancer mutants are hyperactive because they destabilize the double donuts and favor the filament form. We used Surface Plasmon Resonance (SPR) binding assays in which we immobilized full-length BRD3 on the chip at low density and flowed in SPOP as the analyte. We saw a clear gradation in binding affinities with the highest affinity for E47K followed by E78K, intermediate affinities for WT SPOP and D77E, and a lower affinity for V51E (**Table 2**). The off-rates were similar for the different SPOP variants, and the 70-fold difference in *K*_d_ values was largely generated by differences in on-rates (**Table 2**). As a control, we used the prostate cancer mutants W131G and D130E, which carry mutations in the substrate-binding groove of the MATH domain and are loss-of-function mutants ^13^. Their affinities were weaker than that of SPOP V51E, caused mainly by higher off-rates.

**Table 1.** Cryo-EM data collection, refinement and validation statistics.

|  | MATH domains,<br>locally refined | Partial donut,<br>in-phase<br>assembly | Partial<br>donut, out-of-phase<br>assembly | Full double donut |
| --- | --- | --- | --- | --- |
|  | EMDB – 70883 | EMDB –<br>70881 | EMDB –<br>70882 | EMDB – 71281,<br>71282, 71283 |
|  | PDB – 9OUW | PDB – 9OUT | PDB –<br>9OUU | (12mer/13mer/14mer) |
| **Data collection and processing** |  |  |  |  |
| Magnification | 130,000 | 130,000 |  | 79,000 |
| Voltage (kV) | 300 | 300 |  | 200 |
| Electron exposure (e-/Å<sup>2</sup>) | 69.5 | 42.2 |  | 60.5 |
| Defocus range (µm) | -0.8/-2.6 | -0.8/-2.6 |  | -0.8/-2.6 |
| Pixel size (Å) | 0.649 | 0.649 |  | 1.044 |
| Symmetry imposed | C2 | C1 |  | C1 |
| Initial particle images (no.) | 15.4M | 25.5M |  | 2.5M |
| Final particle images (no.) | 391k | 420k | 510k | 15.7/19.7/18.5k |
| Map resolution (Å) | 3.2 | 4.3 | 4.3 | 10.7/9.4/11.3 |
| FSC threshold | 0.143 | 0.143 | 0.143 | 0.143 |
| Map resolution range (Å) | 2.8-7.0 | 3.6-8.7 | 3.6-8.6 | ~8.8-21.8 |
| **Refinement** |  |  |  |  |
| Model resolution (Å) | 3.4 | 4.7 | 4.8 |  |
| FSC threshold | 0.5 | 0.5 | 0.5 |  |
| Map sharpening *B* factor (Å<sup>2</sup>) | -96.1 | -186.5 | -177.2 |  |
| Model composition |  |  |  |  |
| Non-hydrogen atoms | 7706 | 36539 | 35456 |  |
| Protein residues | 956 | 4614 | 4478 |  |
| *B* factors (Å<sup>2</sup>) | 92.6 | 223.4 | 221.0 |  |
| R.M.S. deviations |  |  |  |  |
| Bond lengths (Å) | 0.004 | 0.004 | 0.005 |  |
| Bond angles (°) | 0.862 | 0.902 | 0.805 |  |
| Validation |  |  |  |  |
| MolProbity score | 2.2 | 1.64 | 2.38 |  |
| Clash score | 8.2 | 4.74 | 7.22 |  |
| Ramachandran plot |  |  |  |  |
| Favored (%) | 95.8 | 94.1 | 94.4 |  |
| Allowed (%) | 4 | 5.6 | 5.3 |  |
| Disallowed (%) | 0.2 | 0.2 | 0.3 |  |

**Table 2.** SPR analysis of SPOP variant interactions with BRD3 and Cul3.

| Analyte |  | SPOP |  |  | SPOP |  |  |
| --- | --- | --- | --- | --- | --- | --- | --- |
| Ligand |  | Full-length BRD3 |  |  | Cul3 |  |  |
| SPOP Variant | | $k_{on} (s^{-1}M^{-1})^*$ | $k_{off} (s^{-1})^*$ | $K_d (nM)^*$ | $k_{on} (s^{-1}M^{-1})^*$ | $k_{off} (s^{-1})^*$ | $K_d (nM)^*$ |
| SPOP Variant | E78K | 10397 ± 1324 | 4.0e-4 ± 1.5e-4 | 13 ± 3 | 8694 ± 1562 | 3.4e-4 ± 1.1e-4 | 40 ± 8 |
|  | E47K | 10921 ± 1731 | 1.0e-4 ± 0.3e-4 | 9 ± 3 | 5360 ± 462 | 2.0e-4 ± 0.4e-4 | 43 ± 11 |
|  | WT | 781 ± 134 | 2.8e-4 ± 0.3e-4 | 328 ± 13 | 780 ± 481 | 3.6e-4 ± 1.5e-4 | 673 ± 181 |
|  | D77E | 841 ± 181 | 2.0e-4 ± 0.4e-4 | 228 ± 31 | 2726 ± 1332 | 5.2e-4 ± 2.8e-4 | 186 ± 9 |
|  | V51E | 355 ± 88 | 2.4e-4 ± 0.4e-4 | 681 ± 64 | 3445 ± 2311 | 8.6e-4 ± 5.8e-4 | 657 ± 151 |
|  | D130E | 4457 ± 4354 | 4.5e-3 ± 4.1e-3 | 2352 ± 1368 | 1437 ± 30 | 1.2e-3 ± 0.3e-4 | 831 ± 5 |
|  | W131G | 4485 ± 2269 | 7.5e-3 ± 2.8e-3 | 1728 ± 576 | 3764 ± 2142 | 1.2e-3 ± 8.1e-4 | 309 ± 40 |

\*Mean from at least two replicates ± S.E.M.

Next, we interrogated binding of the series of SPOP variants to the Cul3/Rbx1 complex. We again saw a gradation of *K*_d_ values between SPOP variants, with the highest affinities for E47K and E78K SPOP. Interestingly, WT SPOP bound Cul3 similarly weakly as V51E, suggesting additional complexity in the affinity for Cul3.

Next, we tested whether the series of SPOP variants had biochemical activities in ubiquitination assays that agree with their presumed positions in the hypothetical double donut–filament equilibrium. We used sub-stoichiometric levels of Cul3 to interrogate the effect of the SPOP equilibrium on function. The endometrial cancer mutants were more active than the WT, and the designed mutants were progressively less active, with V51E being the least active (**Fig. 4E**). Stoichiometric Cul3 concentrations resulted in higher initial velocities of ubiquitination as expected while preserving the relative activity differences between variants. Increasing the substrate concentrations enhanced initial velocities even further, again preserving the relative activities of the different mutants.

The most parsimonious explanation for the aggregated data is a model in which SPOP exists in an equilibrium between double–donut and filament states, and the filament state of SPOP is substrate binding-competent and active. Activating mutations enhance the filament population and thereby increase the on-rate for substrates. Given that the bound states should be similar for all variants, the off-rates are also similar. Higher concentrations of Cul3 stimulate the activity by stabilizing the filament state. Upon substrate release, the double donut can reform at a rate determined by the inherent equilibrium of the variant. The double donut thus regulates activity by steric occlusion of the binding site.

### The SPOP double–donut state is evolutionarily conserved and dampens activity in cells

We asked whether SPOP’s ability to form double donuts is conserved among its orthologs. SPOP is conserved throughout the animal kingdom, and most species have a single *SPOP* gene and a close paralog, *SPOPL*, which has an insertion in the BACK domain that prevents formation of oligomers larger than dimers ^24^ (**Fig. S4A**). By contrast, rodents have undergone an expansion in this gene family and possess several SPOP paralogs ^16^. The amino acid sequences of mouse and human SPOP are identical, and mouse SPOP must therefore form double donuts. In addition, we purified 3 of the mouse paralogs and observed that paralog 2 formed double donuts (see the side view in the nsEM image, **Fig. S4B**), while paralogs 1 and 3 formed other types of higher-order assemblies. After the expansion of *SPOP* in rodents, evolution has likely acted independently on the ability of different paralogs to access the double–donut state; nevertheless, in addition to mouse SPOP one of the paralogs forms double donuts suggesting that encoding this property is favorable.

Next, we asked whether the SPOP double donut is also populated in cells. To address this question, we performed in cell ubiquitination assays with the set of variants spanning activating and double donut-stabilizing SPOP variants. E47K and E78K had the highest activity for BRD3 ubiquitination, and the activity of the double donut—stabilizing mutant V51E was lower than that of WT SPOP (**Fig. 5A,B**). Of note, the activity of SPOP D77E was higher than that of the WT although the in vitro data suggested D77E forms double donuts slightly more favorably than WT. Nevertheless, the close resemblance of the degree of cellular and in vitro activities suggests that SPOP double donuts also occur in cells and modulate SPOP activity.

**Figure 5.**
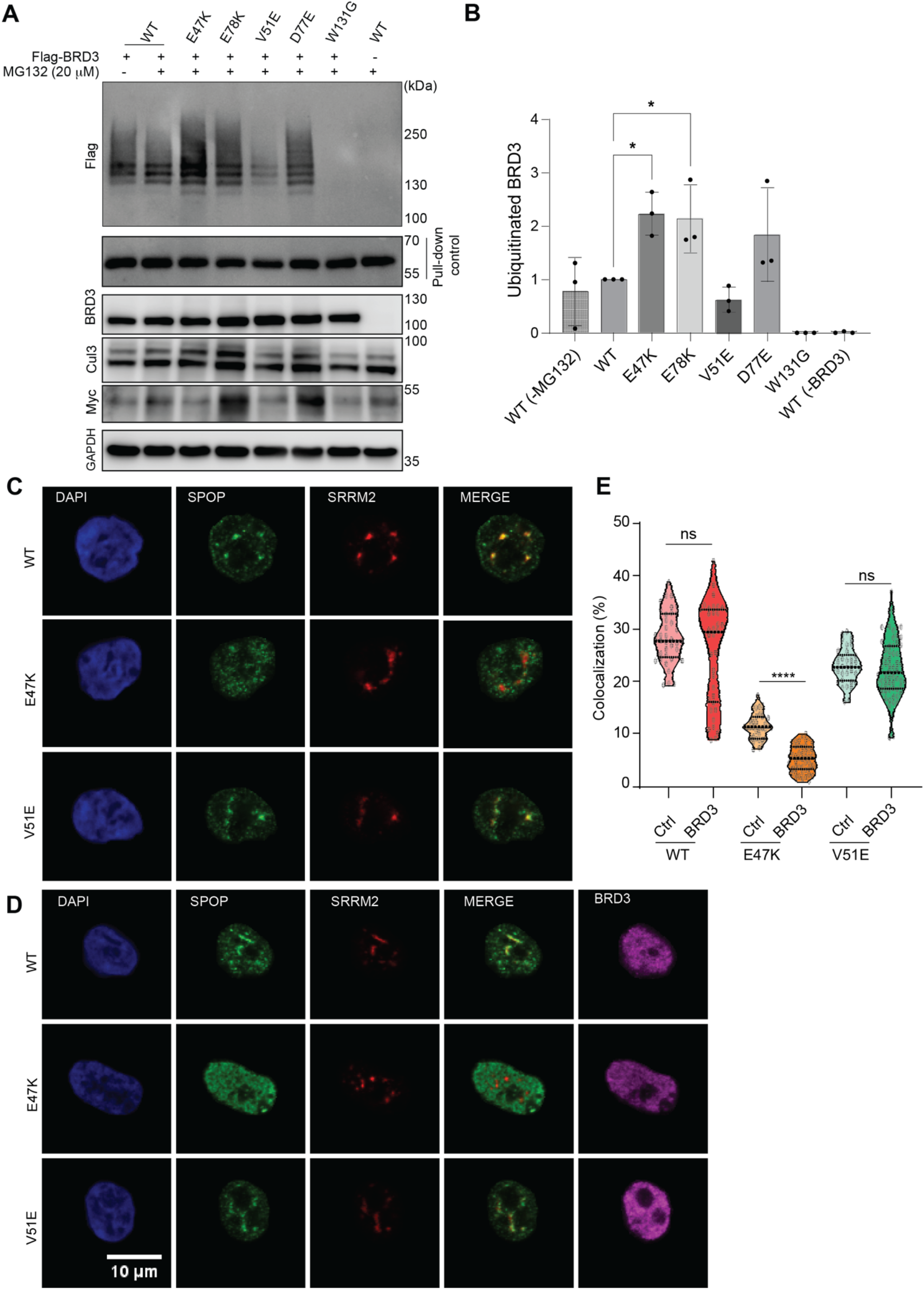
Double donut– vs filament–forming SPOP variants differentially localize to nuclear speckles. **(A)** The SPOP variants along the double donut–filament equilibrium have a wide spectrum of ubiquitination activities towards BRD3 in cells. Representative immunoblot showing ubiquitination in Trex-HEK293 cells co-transfected with Myc-SPOP, Flag-BRD3, His-ubiquitin and Cul3. His-ubiquitin was pulled down using NiNTA sepharose. **(B)** Quantification of BRD3 ubiquitination in the presence of SPOP variants along the double donut–filament equilibrium. The BRD3 signal was quantified and normalized to a control for the effectiveness of pull-down (see Methods). Individual data points (for n=3 experiments) are depicted together with the mean. The error bars indicate the S.E.M. **(C)** Localization of SPOP mutants relative to nuclear speckles. Cells were transfected with the indicated SPOP variants for 24 hrs and stained using anti-Myc (green) and anti-SRRM2 antibodies (the latter to visualize a nuclear speckle component). **(D)** Localization of SPOP variants relative to nuclear speckles upon transient BRD3 expression. Cells were transfected with the indicated Myc-SPOP variants and flag-BRD3 for 24 hrs and stained using anti-Myc (green), anti-flag (magenta) and anti-SRRM2 antibodies. **(E)** Quantification of the fraction of anti-Myc SPOP signal overlapping with SRRM2 foci for SPOP variants with or without transient expression of BRD3.

We used the array of SPOP variants to explore whether there may be differences in double donut and filament localization. A fraction of the SPOP signal typically localizes to nuclear speckles while the rest is found diffuse in the nucleoplasm ^16,20,25^, and this is indeed what we observed for transiently expressed WT SPOP (**Fig. 5C,D**). Endometrial mutants E47K and E78K showed reduced localization to nuclear speckles, while V51E showed a similar localization to nuclear speckles as the WT. This suggests that inactive double donuts preferentially localize to nuclear speckles, while active filaments might be recruited out of nuclear speckles more effectively. When we expressed SPOP variants together with BRD3, we indeed observed that the localization of all variants to nuclear speckles was reduced, but less so in double donut-stabilizing than filament-forming variants (**Fig. 5D,E**), in agreement with their respective abilities to interact with substrates. Nuclear speckles may thus be a storage site for largely inactive SPOP donuts.

### Loss-of-function mutations capitalize on the donut–filament equilibrium

SPOP mutations are found sporadically in cancers other than prostate and endometrial cancer, and if the mutations are not localized to the substrate binding groove, we do not a priori know whether these mutations are activating or deactivating. We interrogated the effects of several additional mutations in the donut-donut interface, including mutations M48I, G128S and R70L found in skin cancer patients (**Fig. 6A**). SPOP M48I and G128S formed rings and had reduced activities in in vitro ubiquitination assays (**Fig. 6B,C**) although they did not affect peptide binding to isolated SPOP MATH domains (**Fig. 6D**) suggesting that they stabilize the double donut state. We next combined the mutations with the double donut–destabilizing E78K mutation. The resulting double mutants had intermediate activities between those of the endometrial cancer and the skin cancer mutants and close to WT activities. SPOP R70L also had lower activity than WT SPOP, but nsEM analysis revealed that it did not form rings but instead double filaments (**Fig. 6C**). Double filaments may have an interface connecting adjacent filaments that is comparable to the donut-donut interface in double donuts. Mutation R70Q was found in intestinal and stomach cancer patients and also gave rise to slightly decreased activity compared to WT. Combination with R78K resulted in intermediate activity.

**Figure 6.**
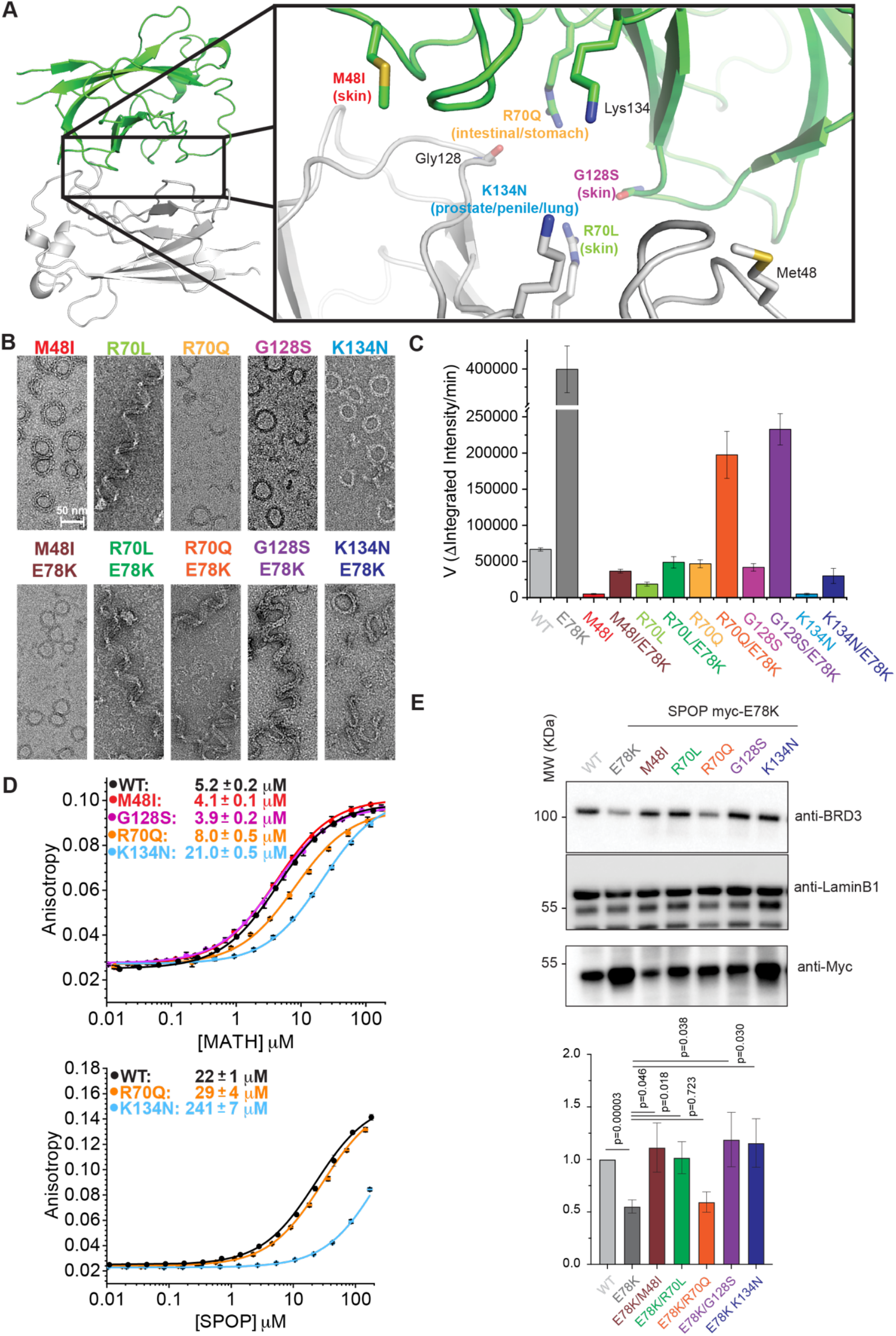
The activities of gain- and loss-of-function SPOP mutants offset each other. **(A)** View of the donut–donut interface where a subset of cancer mutations may stabilize the double donut structure. **(B)** Negative–stain TEM of single mutation variants (top) and double mutation variants, i.e., single mutants in the E78K background (bottom). **(C)** Initial velocities from in vitro ubiquitination assays of single and double variants show that the single variants are deactivating and can be reactivated through combination with the E78K mutation. **(D)** Fluorescence anisotropy binding experiments with a fluorescent BRD3 peptide to determine dissociation constants for binding to either the MATH domain (left) or full-length SPOP (right) WT and variants. The legend reports *K*_D_ values as the mean from triplicate experiments, and the error as the S.D. Wild-type data is the same as in Fig. 1E. **(E)** Quantification of endogenous total BRD3 protein levels in cells normalized to Lamin β1 in the presence of SPOP single and double variants. Top, representative immunoblots show BRD3, Lamin β1 (loading control), and Myc (for SPOP expression verification). Bottom, data represent mean ± S.E.M. from n = 6 independent experiments. Statistical significance was determined using an unpaired t-test.

Typical loss-of-function mutations in prostate cancer patients, such as the frequent mutations to residues Y87, F102, W131 and F133 in the substrate-binding site weaken substrate interactions with these residues directly. By contrast, K134 is located in the donut-donut interface and only borders the substrate-binding groove (**Fig. 6A**). Prostate cancer mutation K134N should therefore not strongly affect the substrate-binding groove. In vitro ubiquitination experiments revealed that K134N SPOP has a lower activity than WT SPOP as expected for prostate cancer mutations (**Fig. 6B)**. nsEM showed that SPOP K134N forms rings (**Fig. 6C**), and we hypothesized that the mutation may reduce substrate binding by shifting the equilibrium towards the inactive double–donut state. Binding of the K134N MATH domain mutant to a substrate peptide was four-fold weaker than of the WT, but full-length SPOP K134N bound 11-fold weaker than full-length WT. These data show that the mutation stabilizes the double–donut state in addition to weakening binding to the substrate-binding groove. Combination with the double donut-destabilizing E78K mutation resulted in close to WT activity (**Fig. 6B**).

We next tested whether this was also the case in cells by assessing steady-state levels of BRD3 in the presence of different SPOP variants. In the presence of E78K SPOP, BRD3 levels were approximately half compared to WT SPOP (**Fig. 6E**) reporting on the high activity elicited by the E78K mutation. Combination of E78K with deactivating mutations M48I, R70Q, G128S and K134N resulted in increased BRD3 levels, similar to those seen in the presence of WT SPOP. Our results reveal that mutations that stabilize double donuts and mutations that destabilize double donuts can be combined to generate intermediate effects in vitro and in cells.

## Discussion

Herein, we discovered that the CRL3–ubiquitin ligase receptor SPOP adopts a double–donut state, a previously unidentified quaternary assembly with a size of up to 2.4 MDa, which dampens its ubiquitination activity. The equilibrium between the inactive double–donut and active filament states (**Fig. 7A**) determines affinities and on-rates for substrates, SPOP ubiquitination activity towards substrates, and the subcellular localization of SPOP. Shifting of the equilibrium towards the filament state via mutations found in patients with endometrial cancer leads to inappropriate activation of SPOP and therefore, we conjecture, promotes oncogenesis (**Fig. 7A**). Unexpectedly, we found that several loss-of-function mutations found in prostate cancer and other types of cancers do not inactivate the substrate binding groove but rather stabilize the double–donut state. We speculate that the canonical prostate cancer mutations of SPOP (e.g., W131G, D130E and K129E) may act simultaneously through perturbation of substrate binding and double donut stabilization, given that they mediate interactions between MATH domains in the double donut (**Fig. 7B**). In support of this view, mutations W131C/R that target one of these sites are found in endometrial cancer patients, likely activating SPOP via the destabilization of double donuts.

**Figure 7.**
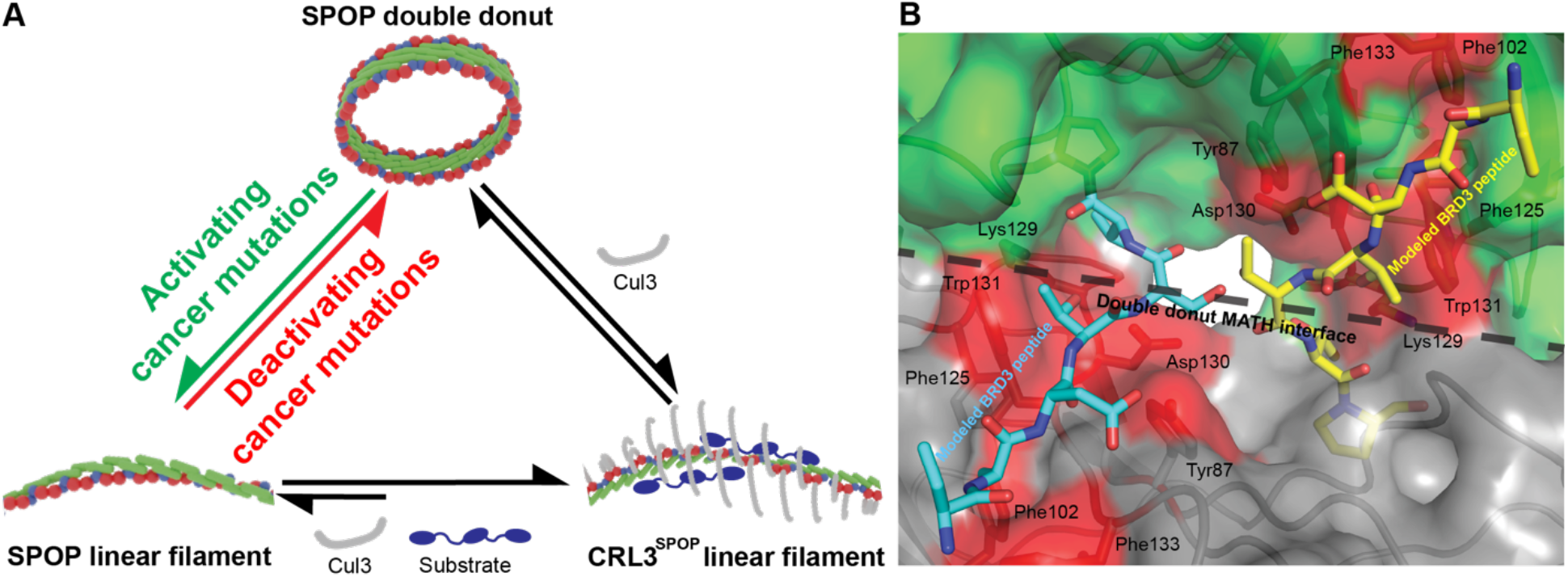
The SPOP double donut–filament equilibrium dictates SPOP activity. **(A)** Schematic of the double donut– filament equilibrium and how it is affected by substrate and Cul3 binding and different types of cancer-associated mutations. Shifting of the equilibrium towards filament or double donut results in hyperactive or hypoactive SPOP, respectively. **(B)** Close-up view of the donut–donut interface with a BRD3 peptide (yellow or cyan carbon atoms, PDB 6I41) superimposed. A subset (residues 129-131) of the canonical prostate cancer mutations (red) also forms interactions across the donut– donut interface, thus potentially contributing to dysregulation through a defect of the binding site and stabilization of the double donut.

The ability of SPOP to form double donuts and the associated regulatory mechanism seems to be evolutionarily conserved. Given that mouse and human SPOP share identical protein sequences, we examined SPOP paralogs that are only found in rodents. Although the expansion of the single human gene to several rodent paralogs implies less evolutionary pressure to maintain a specific regulatory mechanism, one of the mouse paralogs formed donuts, nevertheless (**Fig. S6**).

Regulation of SPOP activity via the double donut–filament equilibrium may explain why SPOP has evolved to form much larger homo-oligomers than other ubiquitin ligase substrate receptors. Only long filaments can circularize and form rings, whose equilibrium with the filament state fine-tunes activity. As opposed to many allosteric enzymes, the activity of SPOP is determined by the equilibrium between two states with large–scale differences in quaternary structure rather than by subtle tuning of side chain packing via filament formation. SPOP regulation is achieved via steric occlusion of the binding site. While other substrate receptors achive this via small oligomeric states ^26^, The SPOP mechanism requires long filaments, and smaller oligomers are unable to engage in this type of steric regulation. The resulting requirement for large, linear homo-oligomers presumably led to co-evolution of substrates with multiple SPOP-binding motifs ^27^, which rendered individual motifs with very weak affinities sufficient ^19,20^. The presence of multiple (if weak) motifs in substrates in turn engenders low off-rates and, consequently, processive poly-ubiquitination ^19^ and phase separation ^20^. We are thus describing a new paradigm of activity regulation that may explain these so far unexplained and unusual features for a ubiquitin ligase substrate receptor.

Regulation of SPOP activity is of key importance given the large number of proto-oncoproteins among its reported substrates including the hormone receptors androgen, estrogen and progesterone receptor; the epigenetic regulators and co-activators BRD2, BRD3 and BRD4 ^14,28,29^; the hormone receptors androgen receptor ^12,30,31^, estrogen receptor ^32,33^ and progesterone receptor ^34^; the co-activator NCOA2 ^35^; the pro-apoptotic factor DAXX ^36^; the SHH regulators Gli2 and Gli3 ^18,37,38^; the proto-oncogenic c-MYC ^39^, and the DNA repair factor 53BP1 ^40^. Clearly, SPOP dysregulation and the attendant dysregulation of the levels of its substrates is risky. The fact that both deactivating and activating mutations can mediate oncogenesis ^14^ further supports this inference. These observations may explain why this elaborate mechanism of SPOP activity regulation has evolved. It provides the ability of both upregulating and dampening SPOP activity. We speculate that the cell may be able to impinge on this equilibrium via post-translational modification to tip the equilibrium towards double donuts or filaments, depending on the cell state and specific signals.

Unexpectedly, we found that mutations that shift the equilibrium towards filament vs double donuts concomitantly alter partitioning of SPOP into nuclear speckles; the filament form preferentially localizes in the nucleoplasm (**Fig. 5**). This may suggest that nuclear speckles house an inactive form of SPOP. However, a small population of E47K SPOP nevertheless localizes to nuclear speckles. Does this fraction represent double donuts, which seem to preferentially partition into nuclear speckles and are therefore stabilized there, or does a small portion of the filament form favored by the endometrial cancer mutant partition into nuclear speckles? Analysis of substrate levels in the presence of WT SPOP *vs* endometrial cancer-associated mutants reported previously ^14^ shows that the endometrial mutants reduce the abundance of a number of nuclear speckle proteins, including SRRM1/2 and FUS as well as numerous splicing factors and RNA-binding proteins common to nuclear speckles. Hence, while WT SPOP may be largely inactive in nuclear speckles, endometrial cancer mutants may retain some activity in nuclear speckles, turn over *de novo* substrates and may therefore not only be hyper-but rather neomorphic.

In conclusion, we report a massive ubiquitin ligase assembly state that functions to sterically fine-tune its activity. We also report a new pathogenic mechanism underlying the inappropriate activation of the ubiquitin ligase via relief of its steric inactivation, and these mutations drive cancer pathogenesis. Furthermore, loss-of-function mutations do not exclusively result from deactivation of the substrate binding site of SPOP but can also be caused by stabilization of the auto-inhibited double–donut state. Given that activating and deactivating mutations can be combined to generate WT-like activities (**Fig. 6**), new therapeutic strategies may be possible if targeted covalent modifications could be used to up- or down-regulate SPOP activity by destabilizing or stabilizing the double–donut state. Our work also has implications for understanding the regulation of filamentous assembly states of other enzymes via large-scale changes in quaternary assembly state and steric occlusion.

## Acknowledgments

We thank Yu-Hua Lo for help with cryo-EM data acquisition and processing of early samples, and Mini José Deepak from the Cell and Tissue Imaging Center and St. Jude Children’s Research Hospital for help with light microscopy. T.M. acknowledges funding by NIH grants R01GM112846 and R01CA301513, and by the American Lebanese Syrian Associated Charities.

