## Supplementary Information for "The equilibrium between two quaternary assembly states determines the activity of SPOP and its cancer mutants"

Supplementary Methods
Supplementary Figures 6
Supplementary References

**Supplementary Methods**
***Plasmids***

The full-length human SPOP gene was cloned into a pCoofy4 vector, which codes for an N-terminal hexa-histidine tag followed by a maltose-binding protein and a 3C protease site, resulting in a construct coding for His-MBP-SPOP. All mutants were created by rolling circle mutagenesis. The full-length mouse SPOP paralogs (paralog 1:Uniprot L7N229, paralog 2:Uniprot Q3UTC4, paralog 3:Uniprot E0CXI6) were cloned into a pDEST-HisMBP[1] vector using Gateway cloning, which contains an N-terminal hexa-histidine tag followed by a TEV site. pDEST-HisMBP was a gift from David Waugh (Addgene plasmid #11085; http://n2t.net/addgene:11086; RRID: Addgene_11085).

The mammalian expression vectors for SPOP WT and mutants were constructed in a manner to give rise to C-terminally Myc-tagged SPOP protein by cloning the coding sequence of mouse SPOP (NCBI: NM_025287.2) into pcDNA4/TO/myc-His (Thermo Fisher Scientific). The expression vector for N-terminally Flag-tagged BRD3 (pEF1α-Flag-BRD3) [2] was generously provided by Joel Mackay (University of Sydney). The expression vector for His-tagged Ubiquitin (pCMV-His-UBC) [3] was generously provided by Wenyi Wei (Harvard Medical School). The vectors for N-terminally HA-tagged RBX1 (pcDNA3-HA2-ROC1, #19897) and for N-terminally Myc-tagged Cullin3 (pcDNA3-myc-CUL3, #19893) were obtained from Addgene. Expression constructs for SPOP mutants were generated by site-directed mutagenesis.

A proximity-based biotin labeling and pull-down technique was employed to investigate the interaction between SPOP and BRD3. To biotinylate the proximal interactors of SPOP, the TurboID biotin ligase coding sequence, kindly provided by Junmin Peng (St. Jude Children's Research Hospital) [4], was cloned into the pcDNA4/TO/myc-SPOP construct.

***Cell lines, cell culture, transfection, and treatments***

T-REX-293 cells were cultured in Dulbecco's modified Eagle's medium (DMEM) supplemented with 10% fetal bovine serum, 1% GlutaMAX, 100 U/mL Penicillin-Streptomycin (Thermo Fisher), and 5 µg/mL Blasticidin S HCl (Santa Cruz) at 37 °C and 5% $CO_2$.

Cells were transfected using Effectene (Qiagen) or PEI-MAX (Polysciences) transfection reagents according to the manufacturer's instructions. To induce SPOP expression, 1mg/ml of tetracycline (Sigma) was added to the culture medium.

***In-cell ubiquitination assays***

T-REX-293 cells were transfected with the following constructs; 12.5 ng of pcDNA4/TO/myc-SPOP, 400 ng of pCMV-His-UBC, 100 ng of pcDNA3-myc-Cul3, 100 ng of pcDNA3-HA2-ROC1, and 100 ng of pEF1α-Flag-BRD3. Tetracycline was added to the culture medium to a concentration of 0.8 µg/mL to induce SPOP-Myc expression. At 24 hrs after transfection, MG132 was added to a final concentration of 20 µM for another 4 hrs. Cells were lysed in 5 mL of buffer A (6 M guanidine-HCl, 0.1 M $Na_2HPO_4/NaH_2PO_4$ pH 8.0, 10 mM imidazole). The lysates were sonicated, cleared, and incubated with 100 µL of Ni-NTA agarose (Qiagen) for a His-tag pull-down. For an internal pull-down control, an equal amount (10 nM) of fluorescently labeled His-tagged *T. maritima* cellobiose-binding protein (tm0031)[5] was added to each sample and further incubated for 30 min. The beads were washed twice with buffer A, twice with A/T buffer composed of one volume of buffer A and three volumes of buffer T (25 mM Tris pH 8.0, 20 mM imidazole), and twice with buffer T. Beads were incubated in 70 µL of SDS-PAGE loading dye containing 300 mM imidazole for 15 min and boiled for 5 min to elute protein.

#### ***SDS-PAGE and Western Blotting***

Protein lysates were denatured in SDS-loading buffer and boiled at 98°C for 5 min. 20-30mg of protein was loaded and subjected to electrophoresis on 4-20% gradient gels (Bio-Rad). Proteins were then transferred to nitrocellulose membranes (Bio-Rad, cat# 1620115) at 90 V for 2 hrs. Next, membranes were blocked with 5% BSA in TBS 0.1% tween-20 (TBST) for 1 hr at RT, followed by over-night incubation with appropriate primary antibody at 4°C. Membranes were washed three times in TBST for 10 min and then incubated with the appropriate HRP-conjugated secondary antibody for 1 hr at RT. After three 10–min washes in TBST and an additional TBS wash, signals were detected using ECL or ECL Select reagents (GE Healthcare). Images were quantified with Image Studio (LI-COR).

#### ***Immunostaining***

T-REX-293 cells were transfected with 12.5 ng of pcDNA4/TO/myc-SPOP with/without 100 ng of pEF1α-Flag-BRD3. Tetracycline was added to the culture medium at a concentration of 0.8 μg/mL to induce SPOP-Myc expression. At 24 h after transfection, cells were fixed with 4% paraformaldehyde, permeabilized with 0.4% Triton-X-100 and then blocked with 3% BSA for 1 hr at RT. They were subsequently, probed with anti-Myc, anti-SRRM2, and anti-Flag antibodies, and appropriate fluorescent secondary antibodies. Images were captured using a Zeiss LSM 780 confocal microscope. To quantitatively assess the colocalization of SPOP with nuclear speckles, immunofluorescence images were analyzed using a combination of image processing and machine learning-based segmentation tools. Multichannel fluorescence images were first separated into individual channels using Fiji (http://fiji.sc). The nuclear speckle signal (SRRM2) channel was then used to train a pixel classification model in Ilastik (http://ilastik.org), enabling accurate identification and segmentation of nuclear speckles. Following model training, nuclear speckle segmentation masks were generated for each corresponding cell. To quantify the degree of SPOP colocalization with nuclear speckles, the SPOP channel was paired with its respective nuclear speckle segmentation mask, and intensity measurements were performed using the MeasureObjectIntensity module in CellProfiler (http://cellprofiler.org). For each cell, the integrated intensity of SPOP overlapping with nuclear speckles was divided by the total nuclear SPOP intensity, representing the fraction of nuclear SPOP localized within nuclear speckles.

#### ***TurboID-based proximity labeling and pull-down assay***

T-REX-293 cells were transfected with pcDNA4/TO/SPOP-myc-TurboID, where the SPOP coding sequence was that of WT or mutants. After 24 hrs, SPOP expression was induced by adding tetracycline (1mg/ml) for 2 hrs. Then, MG132 (20 mM) and biotin (100 mM) were added for an additional 4 hrs. Cells were then harvested and lysed with 8 M urea buffer. Lysates were sonicated and centrifuged at 20,000 x g for 10 min. The resulting supernatant was incubated with 50 μL of Streptavidin agarose beads (Invitrogen) for 16 hrs at RT to enrich for biotinylated proteins. Beads were washed three times with 8 M urea buffer and boiled in 100 μL of SDS–loading buffer for 5 min to elute proteins.

#### ***Protein expression and purification***

His-MBP-SPOP constructs were transformed into BL21-RIPL cells and expressed in autoinduction media. After lysing cells using a microfluidizer in lysis buffer (30 mM imidazole, 1 M NaCl, pH 7.8), the clarified supernatant was mixed with Chelating Sepharose resin, and equilibrated in lysis buffer for 30 min. The resin was applied to a gravity column and washed extensively with lysis buffer, followed by a wash buffer (50 mM imidazole, 500 mM NaCl, pH 7.8). The protein was eluted with elution buffer (250 mM imidazole, 200 mM NaCl, pH 7.8), and 1 mg of 3C protease was added for 1-3 hours. The protein was diluted 3-fold bound to a heparin column, then eluted with a step gradient of NaCl in 20 mM HEPES pH 7.5 with 1 mM dithiothreitol. Mouse paralogs were purified in an identical manner except TEV protease was used in place of 3C protease.

Full-length BRD3 was expressed and purified as previously described.[6] All proteins used for in vitro ubiquitination assays were prepared as previously described.[6] In vitro ubiquitination assays were carried out as previously described except where noted in the text the concentration of components was altered.[6]

#### *Fluorescence anisotropy binding assays*

All fluorescence anisotropy binding assays were carried out in 20 mM Tris pH 7.8, 350 mM NaCl, 0.5 mM DTT. A 12-mer BRD3 peptide that contains the SPOP binding motif (VKRKADTTTDTT) was purchased with a C-terminal PEG linker, followed by 5-carboxyfluorescein. 60 nM peptide was added to serially diluted solutions of SPOP WT or mutants. Fluorescence anisotropy was measured using a CLARIOstar plate reader (BMG LABTECH). Experiments were performed in at least triplicates. Data were fit to a binding equation[7] as previously reported.[8]

#### *Small-angle X-ray scattering*

Small-angle X-ray scattering (SAXS) experiments were performed at the LIX-beamline (16-ID) of the National Synchrotron Light Source II (Upton, NY).[9] Data were collected at a wavelength of 1.0 Å, yielding a scattering angle range of $0.006 < q < 3.2$ Å$^{-1}$, where q is the momentum transfer, defined as $q = 4\pi \sin(\theta)/\lambda$, where λ is the X-ray wavelength and 2θ is the scattering angle. Prior to data collection, SPOP samples were dialyzed into 20 mM HEPES pH 7.5, 400 mM NaCl and 0.5 mM DTT. Samples were loaded into a 1–mm capillary for ten 1–s X-ray exposures. Data were reduced at the beamline using the Python package py4xs and analyzed using Primus.[10,11]

#### *Negative Stain Transmission Electron Microscopy*

SPOP samples (0.01-0.1 mg/mL) were applied to freshly glow–discharged carbon film grids (CF400-CU-50) for 60 s. Solution was wicked away using filter paper and two applications of 4% uranyl acetate solution were used for negative staining. All data were collected on a Thermo Fisher Scientific (TFS) Talos L120C electron microscope, equipped with a 4k x 4k Ceta CMOS camera, at a magnification of 51,000 corresponding to a pixel size of 2.55 Å/px.

#### *Cryo-EM sample preparation and data acquisition*

Non–crosslinked SPOP samples and grids were prepared as previously described.[6] To increase the population of intact SPOP double donuts, which are sensitive to dissociation at the air-water-interface, 1 mg/mL SPOP was crosslinked with 0.0125% glutaraldehyde for 1 hour, and then quenched with 20 mM Tris pH 7.8. For all cryo-EM experiments, 3.0 mL of sample was applied to a freshly glow-discharged grid (C-Flat R1.2/1.3 or C-Flat R2/2, MiTeGen). The grids were blotted for 4.0 s in 100% humidity and plunged into liquid ethane using a Vitrobot mark IV.

To obtain data sets for cross-linked SPOP partial double donuts, the grids were imaged using an TFS Titan Krios G3i with a Gatan Summit K3 electron detector and a BioQuantum Imaging Filter (data collection parameters are shown in **Table 1**). For cross-linked SPOP full double donuts, the grids were imaged using a TFS Talos Arctica with a Gatan Summit K3 electron detector and a BioQuantum Imaging Filter (**Table 1**). The energy filter on both microscopes was set to a slit width of 20 eV. Data collection for non-crosslinked SPOP samples was carried out as previously described (**Table 1**).[6] The exposure of each micrograph was fractioned into 40-70 frames to achieve a dose rate of ~1.0 $e^-/Å^2$/frame and aligned with the MotionCor2[12] software prior to data processing.

#### *Cryo-EM data processing and refinement*

Data processing flowcharts are show in **Fig. S1–S3**. After motion correction and CTF estimation using Cryosparc[13], micrographs were manually curated and filtered. All subsequent data processing was performed in Cryosparc. After particle picking (either template or blob) multiple rounds of 2D classification were used for particle curation. Duplicate particles were removed during the data processing based on a distance cutoff between neighboring particles.

For full double donuts (**Fig. S2**), data were initially processed as partial double donuts (**Fig. S1**). The maps and particle coordinates were then shifted so that the center of mass of the full double donut volume was in the center of a newly extracted box. Duplicate particles were removed prior to and during extensive 2D classification. 3D classification of particles was used after an initial volume was created using ab initio reconstruction. Additional 2D classification was used to further purify the particle sets in a reference–independent manner. These particles were then refined using heterogenous refinement followed by a non-uniform refinement step.[14]

For the locally-refined MATH domains of non-cross linked SPOP, C2 symmetry was applied during non-uniform refinement after an initial heterogeneous refinement.[13] Local refinement with particle subtraction was used to improve the map quality of a hexameric assembly of MATH domains from the partial double donuts.

Molecular models of the partial double donuts and the locally refined MATH domain assemblies were built using Coot and were refined with Phenix.[15,16] The anisotropic resolution of the full double donuts did not permit refinement of models, thus the model presented in **Fig. 3A** was created through superposition of the refined partial double donuts into the map volume using ChimeraX.[17] No further refinement or modelling was carried out.

#### *SPR analysis of SPOP interactions*

Surface plasmon resonance (SPR) experiments were performed using a Biacore T200 instrument (Cytiva) equipped with a CM5 sensor chip to characterize the binding kinetics of His-tagged protein interactions. Anti-histidine antibodies were immobilized onto both the reference and active flow cells of the sensor chip using standard amine coupling chemistry with N-hydroxysuccinimide (NHS) and 1-ethyl-3-(3-dimethylaminopropyl)carbodiimide hydrochloride (EDC), following the protocol provided in the His Capture Kit (Cytiva). Residual active esters were quenched with 1 M ethanolamine-HCl (pH 8.5). After coupling, the sensor chip was equilibrated with 20 mM HEPES pH 7.5, 200 mM NaCl, 0.5 mM DTT for 2 hours at 25°C. The same buffer was used throughout the experiment as the running buffer and for preparing analyte dilutions.

His-tagged Cul3 or BRD3 proteins (each with an approximate molecular weight of 85 kDa) were captured onto the active flow cell by injection for 120 seconds at 10 µL/min, resulting in immobilization levels of approximately 200 response units (RUs). The reference flow cell contained only the immobilized anti-histidine antibody and served to correct for nonspecific binding and refractive index changes.

Binding kinetics were evaluated using a multicycle kinetic analysis format. Analytes were injected at 50 μL/min across a concentration series ranging from 0.015 μM to 2 μM (eight concentrations in 2-fold serial dilutions), with each cycle consisting of a 120-second association phase and a 600-second dissociation phase. Buffer-only injections were included between sample injections to facilitate double referencing.

The sensor surface was regenerated between cycles using sequential injections of 1 M NaCl followed by 10 mM glycine-HCl (pH 2.0), ensuring complete removal of bound analyte prior to recapture.

Data analysis was performed using the Biacore T200 evaluation software. Sensorgrams were processed by subtracting both the reference cell and buffer blank responses. Kinetic constants were obtained by global fitting to a 1:1 Langmuir binding model, from which the association ($k_a$) and dissociation ($k_d$) rate constants were derived. The equilibrium dissociation constant ($K_d$) was calculated as $k_d/k_a$. Reported $K_d$ values represent the mean from at least three independent experimental replicates.

### Supplementary Figures

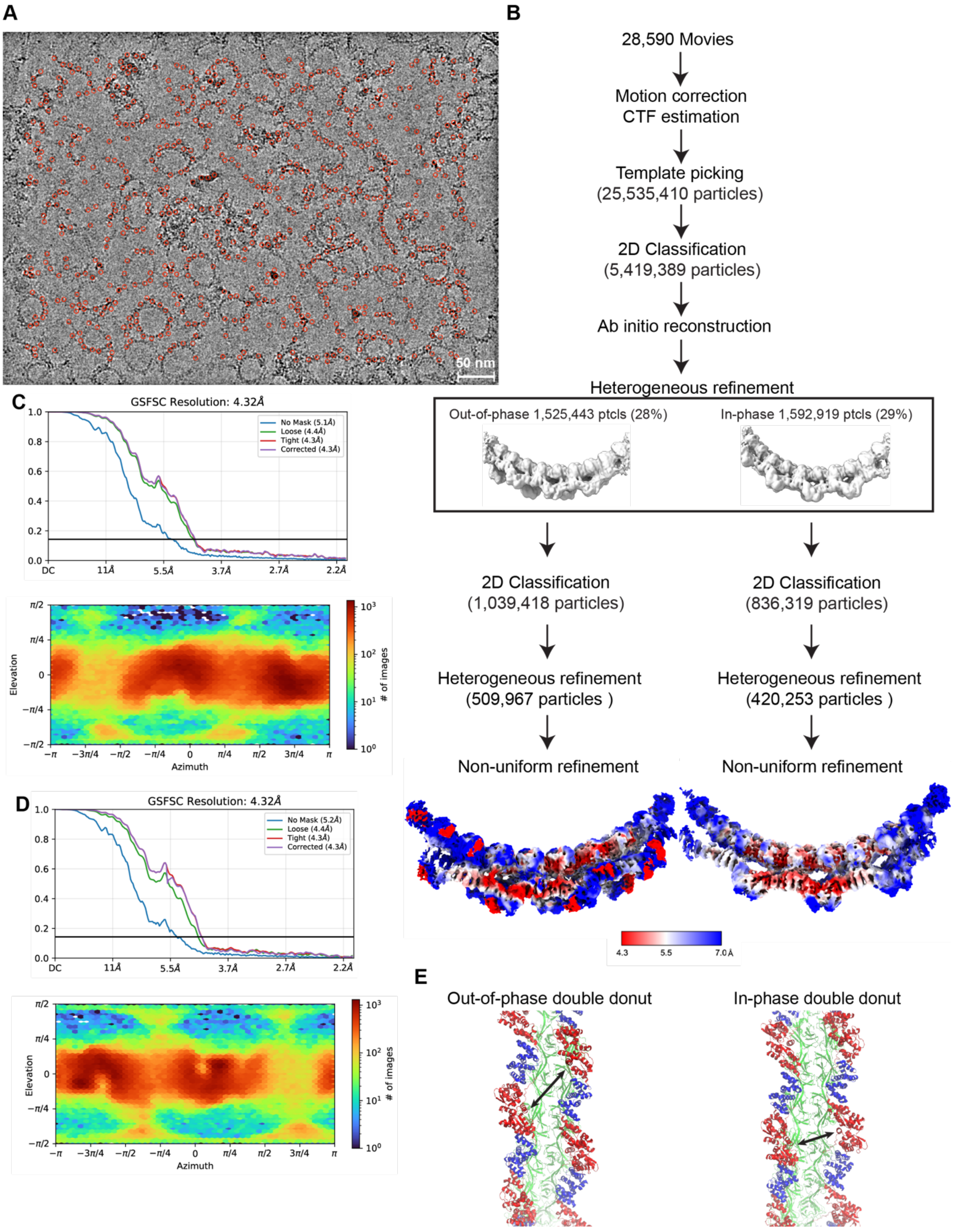


**Figure S1. Cryo-EM data processing and validation of partial double donut assemblies.**

**(A)** Representative micrograph with picked particles indicated by red circles. **(B)** Data processing strategy. Bottom, refined maps of out-of-phase (left) and in-phase (right) partial double donut assemblies colored by resolution. **(C)** Fourier Shell Correlation plots (top) and viewing direction distribution plot of partial double donut, out-of-phase assembly. **(D)** Fourier Shell Correlation plots (top) and viewing direction distribution plot of partial double donut, cis assembly. **(E)** Side view of the refined partial double donut assemblies showing the shifted register of BTB/BACK domains in in-phase *vs* out-of-phase assemblies.

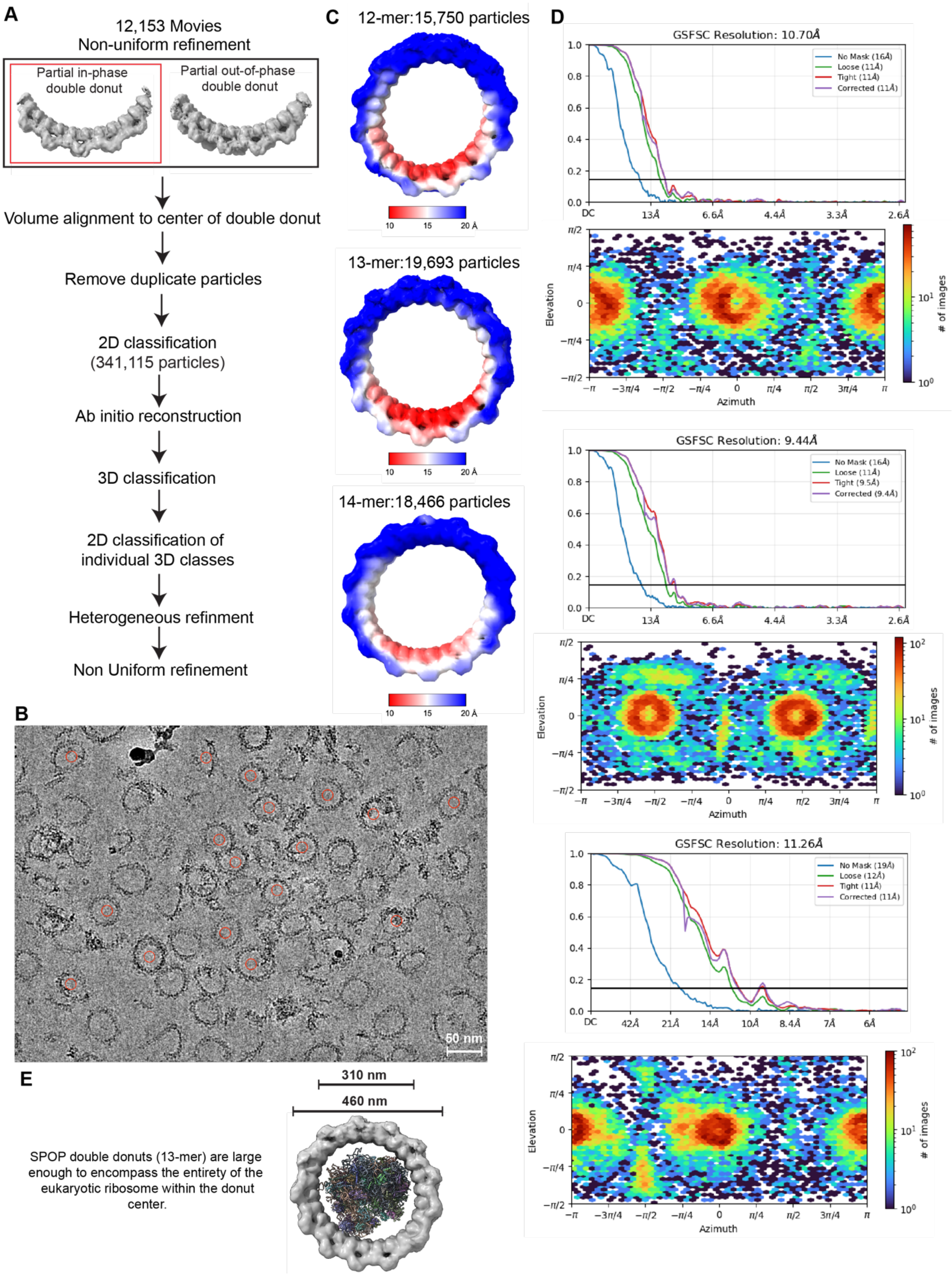


**Figure S2. Cryo-EM data processing and validation of full double donut assemblies.**

**(A)** Full donuts were processed initially in a manner identical to the strategy in **Fig. S2B**. These particles were then further processed after volume alignment to the center of the donut particles. **(B)** Representative micrograph with picked particles indicated by red circles. **(C)** Cryo-EM maps and particle counts used in the generation of maps for the 12-mer, 13-mer and 14-mer double donut assemblies. Maps are colored by resolution. **(D)** Fourier Shell Correlation plots (top) and viewing direction distribution plot of 12-mer in-phase double donut (top), 13-mer in-phase double donut (middle) and 14-mer in-phase double donut (bottom). **(E)** The size of the double donut assemblies is larger than the eukaryotic ribosome (PDB code 6GZ5).

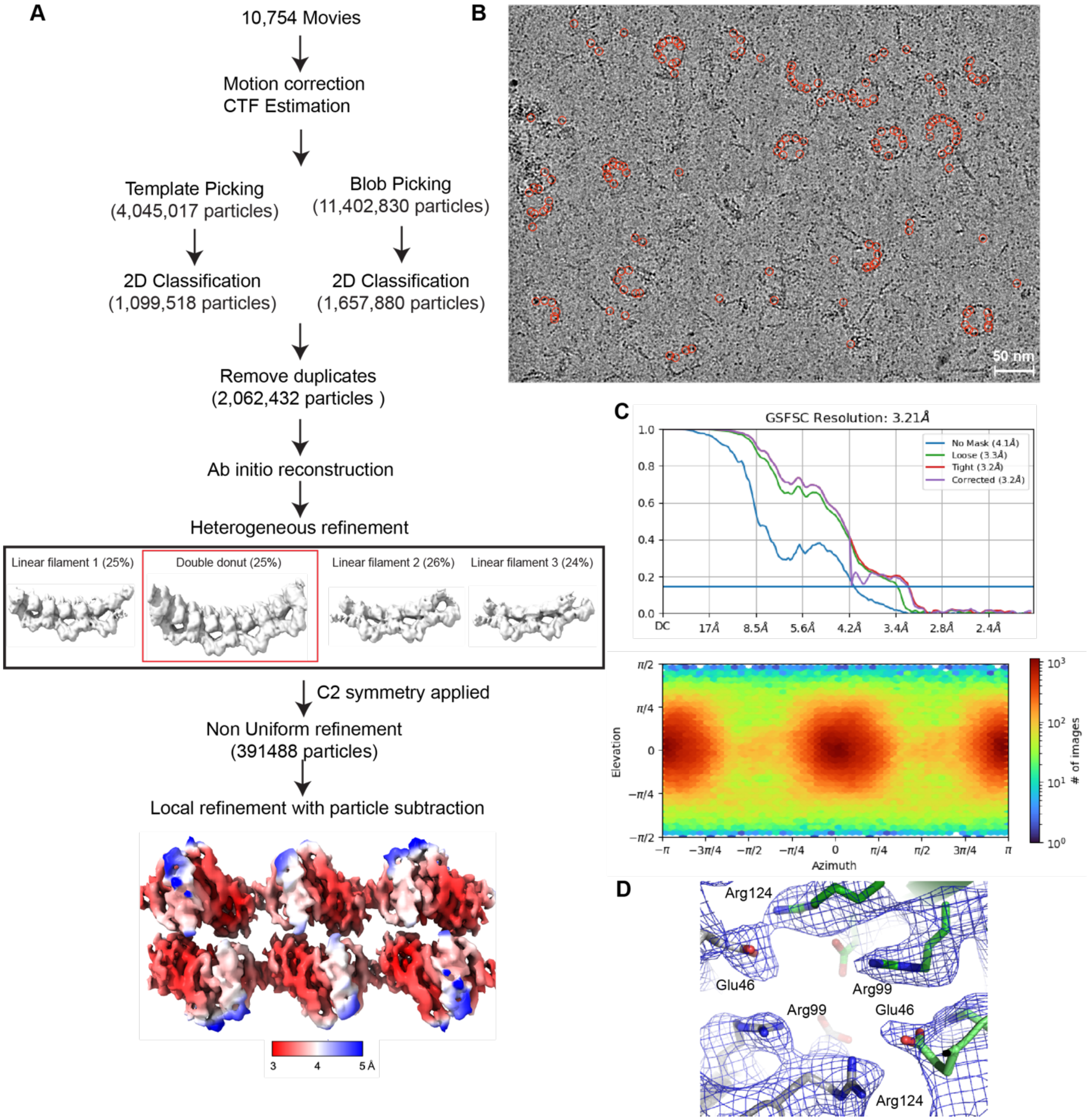


**Figure S3. Cryo-EM data processing and validation of non-cross-linked locally refined MATH domains.**
**(A)** Data processing strategy. Bottom, refined maps of locally refined MATH domains colored by resolution. **(B)** Representative micrograph with picked particle indicated by red circles. **(C)** Fourier Shell Correlation plots (top) and viewing direction distribution plot of locally refined MATH domains. **(D)** Cryo-EM maps of the double–donut MATH–MATH interface showing resolution of map for individual side chains.

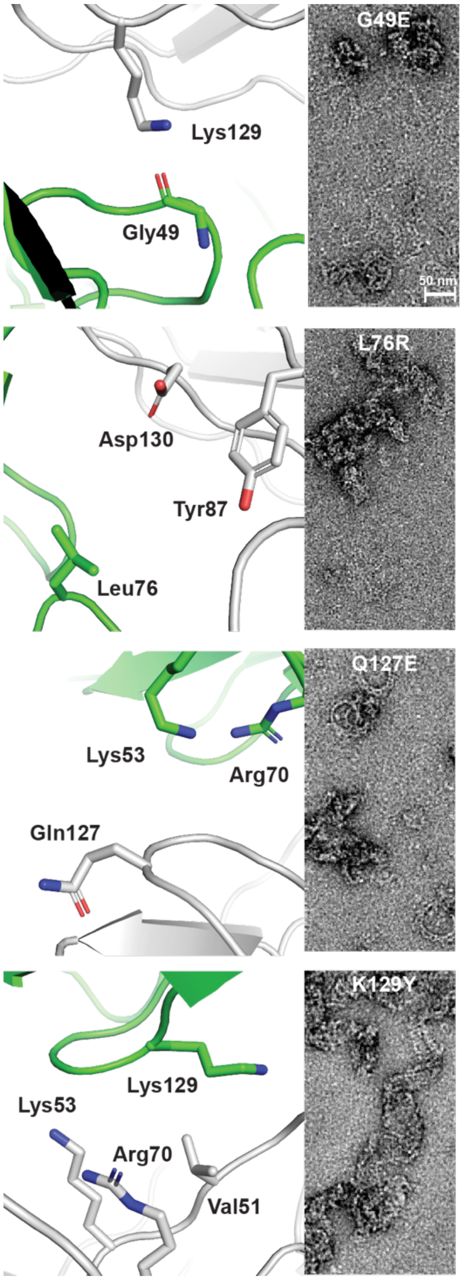


**Figure S4. Designed mutants that lead to the disruption of the ring state.**
Left, the high-resolution structure of the MATH assemblies across the donut–donut interface was used to identify candidate mutations (left panels). The mutants were subjected to negative–stain TEM (right panels), and this subset of mutants disrupted the ring state.

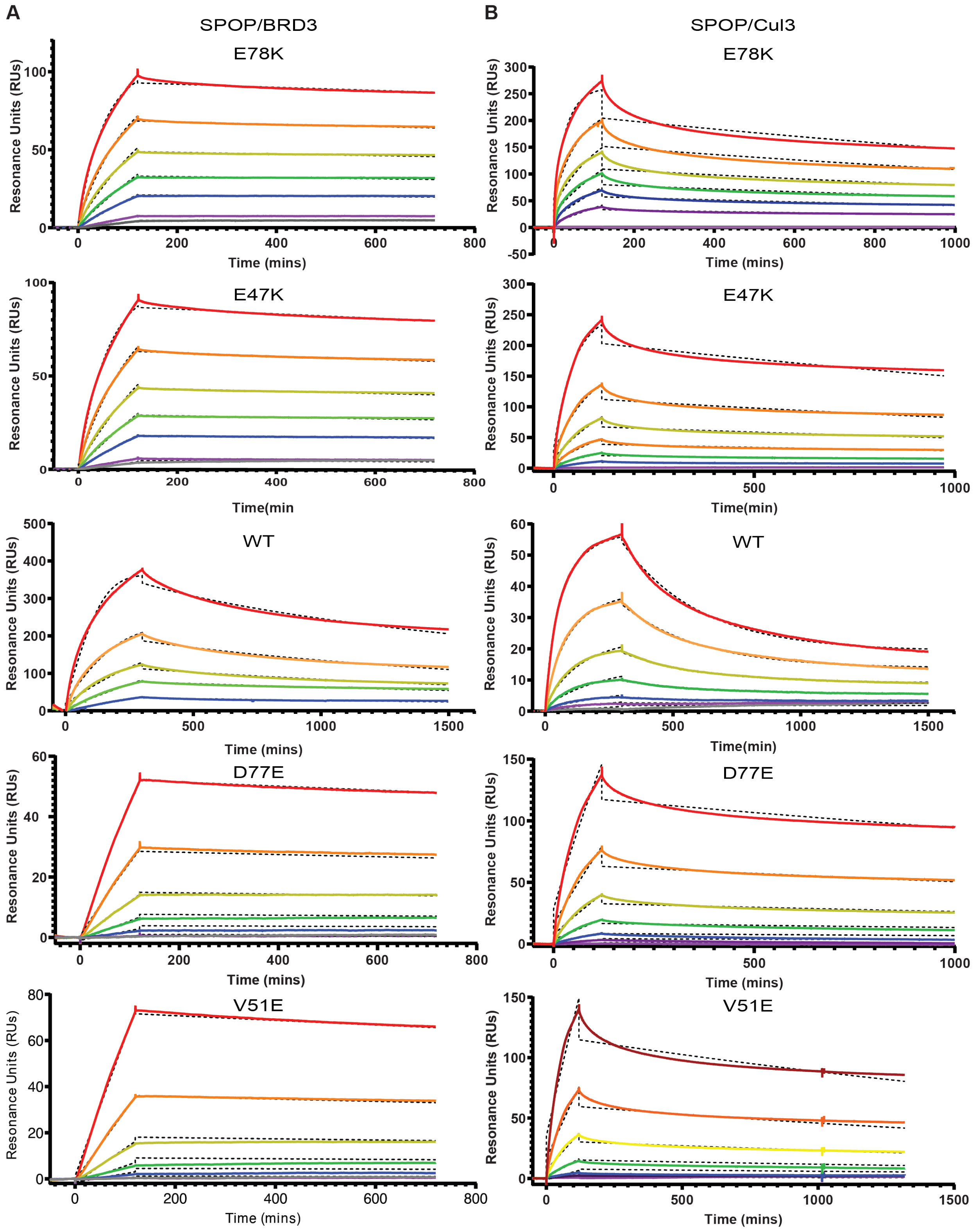

A
SPOP/BRD3
B
SPOP/Cul3
E78K
E47K
WT
D77E
V51E
Resonance Units (RUs)
Time (mins)
Time(min
Time(min)

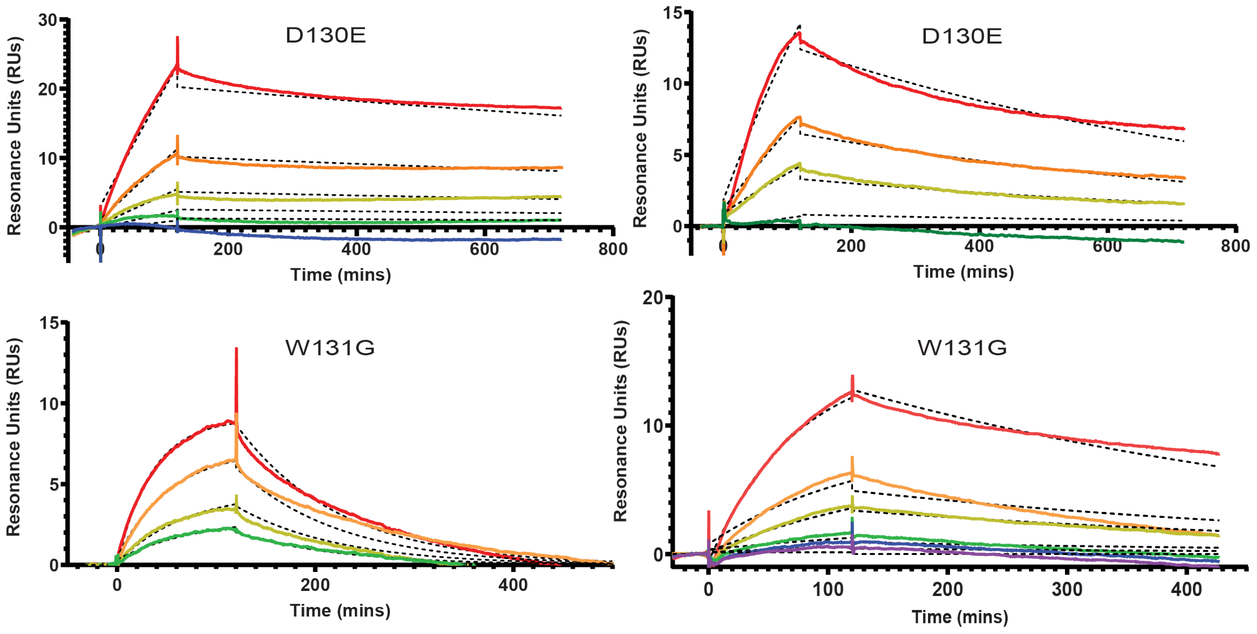


**Figure S5. Representative plots of SPR data, related to Table 2. (A)** SPR traces (solid lines) where BRD3 is the ligand and SPOP variants are the analyte at increasing concentrations from cold to warm colors. Dashed lines are a fit to the data. **(B)** SPR traces (solid lines) where Cul3 is the ligand and SPOP variants are the analyte. Dashed lines are a fit to the data.

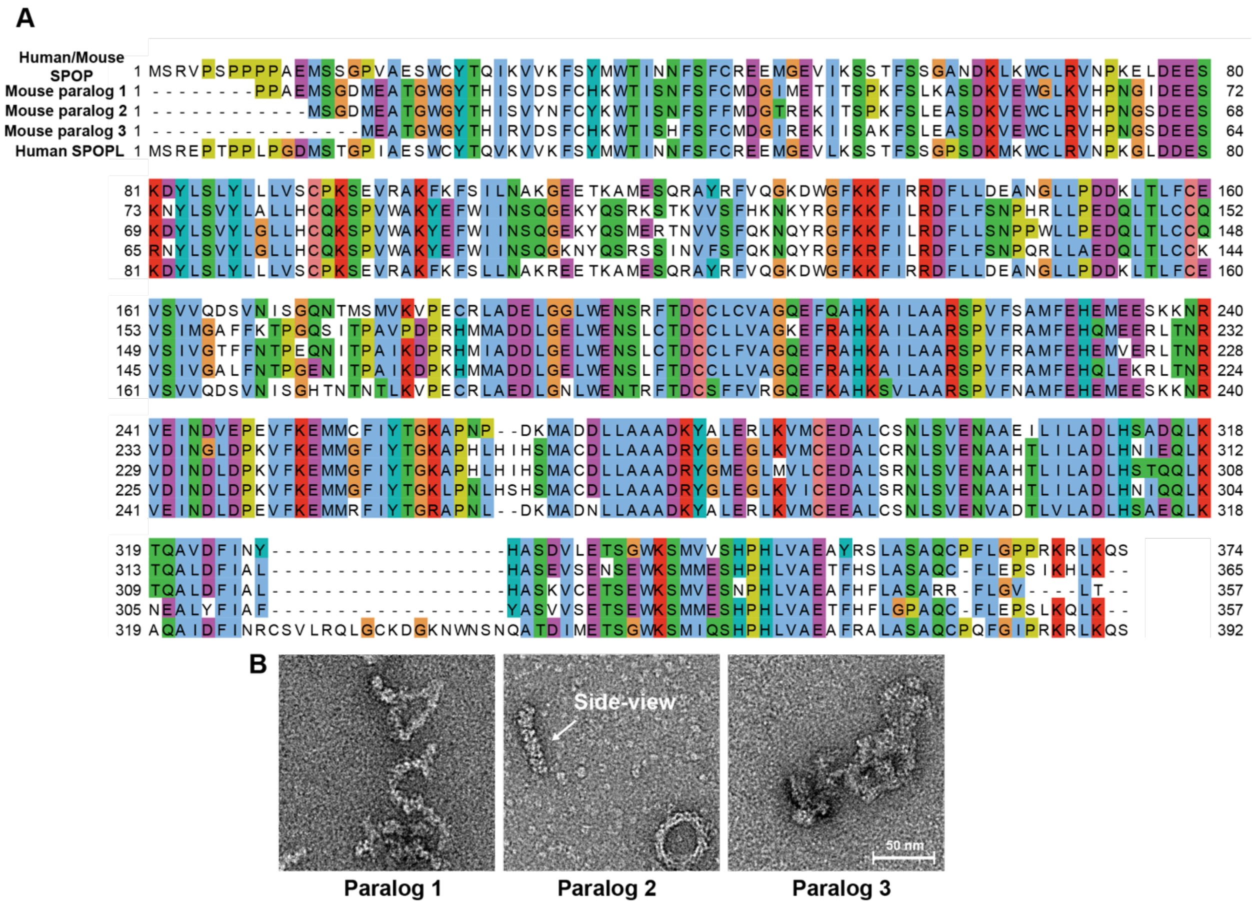


**Figure S6. The ability of SPOP to adopt double donut assemblies is conserved in a mouse paralog, related to Figure 5. (A)** Amino acid sequence alignment of three additional paralogs of SPOP found in the mouse genome together with human/mouse SPOP (which have identical protein sequences) and human SPOPL. **(B)** Negative–stain TEM of the three mouse paralogs shows double donut assemblies are adopted in one of the three.
